# Electroencephalographic Biomarkers of Relaxation: A Systematic Review and Meta-analysis

**DOI:** 10.1101/2024.03.27.586444

**Authors:** Kairi Sugimoto, Hideaki Kurashiki, Yuting Xu, Mitsuaki Takemi, Kaoru Amano

## Abstract

The human electroencephalogram (EEG) is composed of synchronous oscillations within characteristic frequency bands, including 8–13 Hz alpha oscillations that often appear during relaxation. Relaxation is critical for physical and mental health, but the extent to which various EEG components reflect relaxation remains uncertain. This systematic review and meta-analysis investigated associations between EEG components and concurrently measured reference indices of relaxation in healthy adults. A comprehensive database search and screening using preset criteria identified 38 studies involving 1,120 participants published between January 1940 and January 2022 for qualitative synthesis. These studies used various reference relaxation measures such as electrocardiographic indices related to parasympathetic nervous system activity and introspective indices obtained through questionnaires. Risks of bias were evaluated following the risk of bias assessment tool for nonrandomized studies. A meta-analysis of 23 studies using a random-effects model revealed positive correlations between relaxation index and the power of alpha oscillations at central channels (Fisher’s *z*-transformed correlation coefficient and [95% confidence interval]: 0.23 [0.10–0.36]) and frontal channels (0.16 [0.02–0.31]). The correlation was significantly higher for the central channel compared with other channels. No significant correlations were detected between relaxation indices and other EEG frequencies or channels. The causal relationships between relaxation and alpha power at central and frontal channels warrant further study.

## Introduction

Relaxation refers to a state of body and mind free from tension, stress, and anxiety. Practicing relaxation helps reduce physiological and psychological stress symptoms and can foster a sense of well-being (Sianoja et al., 2018). Across different cultures, relaxation methods have been developed to enhance overall health and well-being. Notably, meditation and yoga have long attracted the attention of clinicians and therapists as interventions that induce relaxation for improved physical and mental health (Gillani & Smith, 2001; Jain et al., 2007; Manzoni et al., 2008; Vempati & Telles, 2002). More recent methods include pleasant and calming virtual environments experienced through immersive head-mounted displays (Riches et al., 2021; Velana et al., 2022) and neurofeedback techniques to achieve relaxation utilizing electroencephalogram (EEG) measurements (Biswas & Ray, 2019; Grosselin et al., 2021).

Precise assessment of the state of relaxation is pivotal for appraising the effectiveness of relaxation interventions and maximizing health outcomes. The classical view of relaxation posits that it is accompanied by a distinctive pattern of autonomic nervous system (ANS) activity (Lucini et al., 1997). Measures of parasympathetic nervous system activity, such as electrocardiographic (ECG)-related indices (Hernando et al., 2018; Tan et al., 2015), respiratory activity (Telles et al., 2000), blood pressure (Santaella et al., 2006), skin conductance (Khalfa et al., 2002), and saliva composition (Knight & Rickard, 2001), have been employed to evaluate the state of relaxation. Among these measures, ECG-related indices, notably heart rate variability (HRV), are the most widely used for both research and commercial applications because of their reliability in reflecting ANS activity and the ease of continuous measurement by wearable devices (Hernando et al., 2018; Malik et al., 1996; Sztajzel, 2004; Zhang et al., 2020).

In addition to these physiological measures, EEG alpha oscillations, synchronous voltage oscillations occurring within the frequency range of 8 to 13 Hz, are often assumed to represent a relaxed state, particularly by the general public (Cloud, 2011; TIME, 1971). This idea may stem from the typical predominance of alpha oscillations in people at rest with eyes closed (Adrian & Matthews, 1934; Berger, 1929; Goldman et al., 2002) or fully awake but not doing or thinking about anything in particular (Berger, 1929). However, the empirical evidence linking specific cortical oscillatory activity measures to relaxation remains inconclusive. This ambiguity is due primarily to the complex functional nature of resting alpha oscillations, which are influenced not only by the state of relaxation but also by unrelated factors such as sleepiness (Putilov et al., 2017; Strijkstra et al., 2003), attention and inhibition (Chang & Huang, 2012; Foxe & Snyder, 2011; Kelly et al., 2006; Klimesch, 2012), mental fatigue (Li et al., 2020; Tran et al., 2020), and emotional state (Allen et al., 2018).

Furthermore, relaxation may be related to EEG frequency bands beyond alpha. For instance, mindfulness and meditation, which are frequently employed relaxation interventions, have been reported to influence theta (4–8 Hz), beta (13–30 Hz), and gamma (>30 Hz) oscillations in addition to alpha oscillations depending on the specific practice (Chan et al., 2008; Hinterberger et al., 2014; Lee et al., 2018; Lomas et al., 2015; Sinha et al., 2020). Meditation has also been reported to modulate theta and alpha oscillations (Lomas et al., 2015; Rodriguez-Larios et al., 2020). Moreover, the relationships between EEG oscillations and physiological functions are influenced by factors such as age (Dustman et al., 1985), sex (Carrier et al., 2001), physical activity habits (Lardon & Polich, 1996), and meditation experience (Lomas et al., 2015). This complexity provides a compelling rationale to comprehensively examine the relationship between neural oscillations and relaxation across a wider frequency range and across studies using a variety of distinct relaxation measures and techniques.

In this study, we conducted a systematic review and meta-analysis of the relationship between EEG frequency band and the state of relaxation in healthy adults. While a few studies have focused on a specific type of relaxation such as mindfulness (Lomas et al., 2015), transcendent states (Wahbeh et al., 2018), breath control (Zaccaro et al., 2018), essential oil inhalation (Komini et al., 2023), indoor plants (Han et al., 2022), and urban built and natural environment (Bolouki, 2023), or a specific population such as depression and anxiety patients (Thibodeau et al., 2006), children and adolescents with long-term physical conditions (Thabrew et al., 2018), and people with epilepsy and intellectual disabilities (Jackson et al., 2015), no systematic review or meta-analysis has examined the general relationship between EEG pattern and relaxation among healthy adults. Revealing universally applicable relationships among healthy adults is expected to have broad utility in diverse domains related to relaxation and contribute to the responsible utilization of brain information for human benefit. Moreover, we planned to conduct subgroup meta-analyses considering EEG electrode positions based on the international 10–20 system and reference indices reflecting relaxation states such as ECG measures of ANS activity and introspective reports. The miniaturization and reduced cost of EEG measurement devices has also allowed for the accurate assessment of physiological states in real-world settings, so these findings are relevant across diverse fields related to relaxation, from basic research to consumer product development.

## Methods

The systematic review and meta-analysis were conducted following Chapter 4 of the Minds Manual for Clinical Practice Guideline Development 2020 version 3.0 (Minds Manual Developing Committee, 2021). The review question was formulated using a modified version of PICO (Population-Intervention-Comparators-Outcome) for diagnostic test accuracy. The PICO in this study is as follows: Population, healthy human adults (18–65 years); Intervention, EEG/MEG measurement; Comparator, primary indicator of relaxation other than EEG (reference index); Outcome, correlation between EEG/MEG and reference relaxation indices. The authors had expertise in systems neuroscience, human neuroimaging, or cognitive psychology.

### Literature Search

Two authors (HK and KS) independently performed a literature search on January 14, 2022, using the following databases: PubMed, Scopus, Web of Science, JDreamⅢ, and Ichu-shi. JDreamⅢ and Ichu-shi were used to access scientific and medical studies written in Japanese, respectively. The search string of keywords was constructed to include the measurement system (electroencephalogra* OR eeg OR magnetoencephalogra* OR meg), targeted outcome (relax OR relaxation), and population (child* OR infant* OR kid OR kids OR animal* OR monkey* OR rodent* OR rat OR rats OR cat OR cats). The measurement and outcome parts were combined using “AND,” and the population part was used to exclude studies if those words were included in the title or keywords. The inclusion of the comparator component was considered during the screening processes. Both English and Japanese terms were used for the Japanese databases. The language restriction for all databases was set to English or Japanese. For databases other than PubMed, filtering was applied to include only original articles. The exact search queries for each database can be found at <u>the OSF repository</u>.

### Study Selection

After removing duplicates retrieved from the literature search of all databases, studies were selected in two screening phases according to PRISMA recommendations (Page et al., 2021). In the first screen, all studies were divided into three sets, and two authors ([HK and KS], [KS and YX], or [HK and YX]) independently reviewed the titles and abstracts for eligibility according to the following inclusion criteria (1)–(5): (1) written in English or Japanese; (2) published as an “original article” in a scholarly peer-reviewed journal; (3) participants were healthy adults aged 18–65; (4) EEG/MEG measurements were conducted in the awake state; and (5) reference relaxation indices (other than EEG/MEG) were measured during the awake state. Studies for which authors disagreed about relevance/inclusion were retained for the second screening.

During the second screening, three authors (HK, KS, and YX) independently conducted full-text assessments according to inclusion criteria (6) and (7) in addition to the aforementioned criteria (1)–(5): (6) EEG/MEG and reference indices were measured at the same time or in succession and (7) correlation coefficients between EEG/MEG and reference indices were reported, or raw data for the calculation of correlation coefficients were available. Two authors (KA and MT) participated in the screening process in cases of disagreement. The five authors engaged in discussions until a consensus was reached regarding study inclusion.

Studies that met the following exclusion criteria (8)–(10) were then removed from the meta-analysis: (8) judged to have a high risk of bias in more than one domain; (9) judged to have unclear or high indirectness in more than one domain; and (10) the directions of numerical changes in reference indices were unclear. Following the application of these criteria, studies were also excluded from the meta-analysis if they utilized EEG indices which were only employed in fewer than two studies employing the respective indices were identified (e.g., microstate, beta/alpha ratio), as this would not support robust variance estimation. Consequently, only studies reporting EEG indices related to oscillation power/amplitude were included in the meta-analysis. If there was any doubt about the data, we contacted the corresponding author to request the relevant information. If no reply was received, the study was also excluded.

All types of interventional study designs (randomized controlled trials, nonrandomized studies of interventions, and uncontrolled studies) were included in the meta-analysis because the risk of bias in the correlation between EEG/MEG and reference indices was deemed to be consistent across these study designs. As this meta-analysis focused on evaluating correlations rather than intervention outcomes, potential differences in the risk of bias among different study designs were considered negligible, and effect sizes from all study designs were included. This approach effectively treated the meta-analysis as if virtually all included studies were NRSIs.

### Qualitative Assessment

Qualitative assessment of individual studies was conducted independently by two authors (HK and KS). Only studies that successfully passed the second screening were included in the qualitative assessment. The following data were first extracted: study population (including sample size and participant demographics), information related to the EEG indices (such as types of feature components, electrode positions, timing of measurements, cumulative period used for calculating a single EEG index), types of reference indices, and other variables such as year of publication, study design, and intervention and control conditions if applicable. As mentioned, we contacted the corresponding author to request the relevant information if there was any doubt about the data. The extracted data are summarized in the Tables, and a subset is presented in the Figures.

### Risk of Bias Assessments

Risk of bias assessments for individual studies that passed the second screening were performed independently by two authors (HK and KS) according to the risk of bias assessment tool for nonrandomized studies (Kim et al., 2013). The risk of bias was rated as low, unclear, or high for six domains: selection bias, performance bias, detection bias, attrition bias, reporting bias, and other possible sources of biases. Disagreements were resolved through discussions among four authors (KA, HK, KS, and MT).

Selection bias was rated by considering the allocation method, baseline imbalance, and confounding factors. Performance bias was assessed based on three criteria: blinding of measurements, consistency of measurement indices, and types of measurement indices. The consistency of measurement indices refers to the utilization of identical indices for all participants. The types of measurement indices were appraised according to whether they were introspective measures (e.g., from questionnaires) and thus susceptible to interviewer bias or recall bias (Kim et al., 2013). Detection bias is associated with researcher blinding in outcome assessments. The detection bias was considered low irrespective of blinding conditions if we were able to calculate correlation coefficients from the reported raw data. Attrition bias was determined by the proportion of participants excluded and the reasons for exclusion. Reporting bias was verified by the consistency between the reported and preregistered outcomes. Bias was considered low when correlation coefficients could be calculated from the reported raw data. Other possible sources of bias were judged according to sample size setting and conflict of interest (COI) disclosure.

### Indirectness Assessment

The assessment of indirectness for individual studies that passed the second screening was conducted independently by two authors (HK and KS) across four domains: participants, EEG/MEG indices, targeted outcomes, and physiological state/relaxation indices. Indirectness related to participants, EEG/MEG indices, and targeted outcomes was classified as either low or unclear. If there was no description of participant selection, EEG/MEG recording methods, or calculation procedure of EEG/MEG indices, indirectness in the respective domain was considered unclear. Similarly, if correlation coefficients were calculated from sparse discrete measures, such as a questionnaire with items rated on four levels or less, indirectness regarding targeted outcome was considered unclear.

Indirectness regarding the physiological state captured by the EEG/MEG index and the relaxed state measured by reference indices was classified as either low, high, or unclear. If the EEG/MEG measurement was not conducted during, right before, or right after the measurement of reference indices, increasing the probability that the physiological state captured by the EEG/MEG index does not correspond to the relaxed state measured by reference index/indices, indirectness was assumed to be high. If there was no description of the temporal relationship between EEG and reference index measurements, indirectness was considered unclear.

### Meta-analysis

#### Data Extraction

Pearson’s and Spearman’s correlation coefficients between EEG/MEG indices and reference indices for the relaxation state were calculated as outcome variables. They were independently extracted by two authors (HK and KS), and an experiment in each paper was regarded as a single trial. For studies that did not report correlation coefficients but provided raw data or other statistics and met all other inclusion criteria set for the second screening, two authors (HK and KS) calculated the correlation coefficients. Furthermore, because some reference indices increase with higher relaxation levels while others decrease (Table 1), we adjusted the sign of the correlation coefficients such that a larger positive *r*-value always indicates that an improved EEG/MEG index reflects a greater state of relaxation.

**Table 1.**
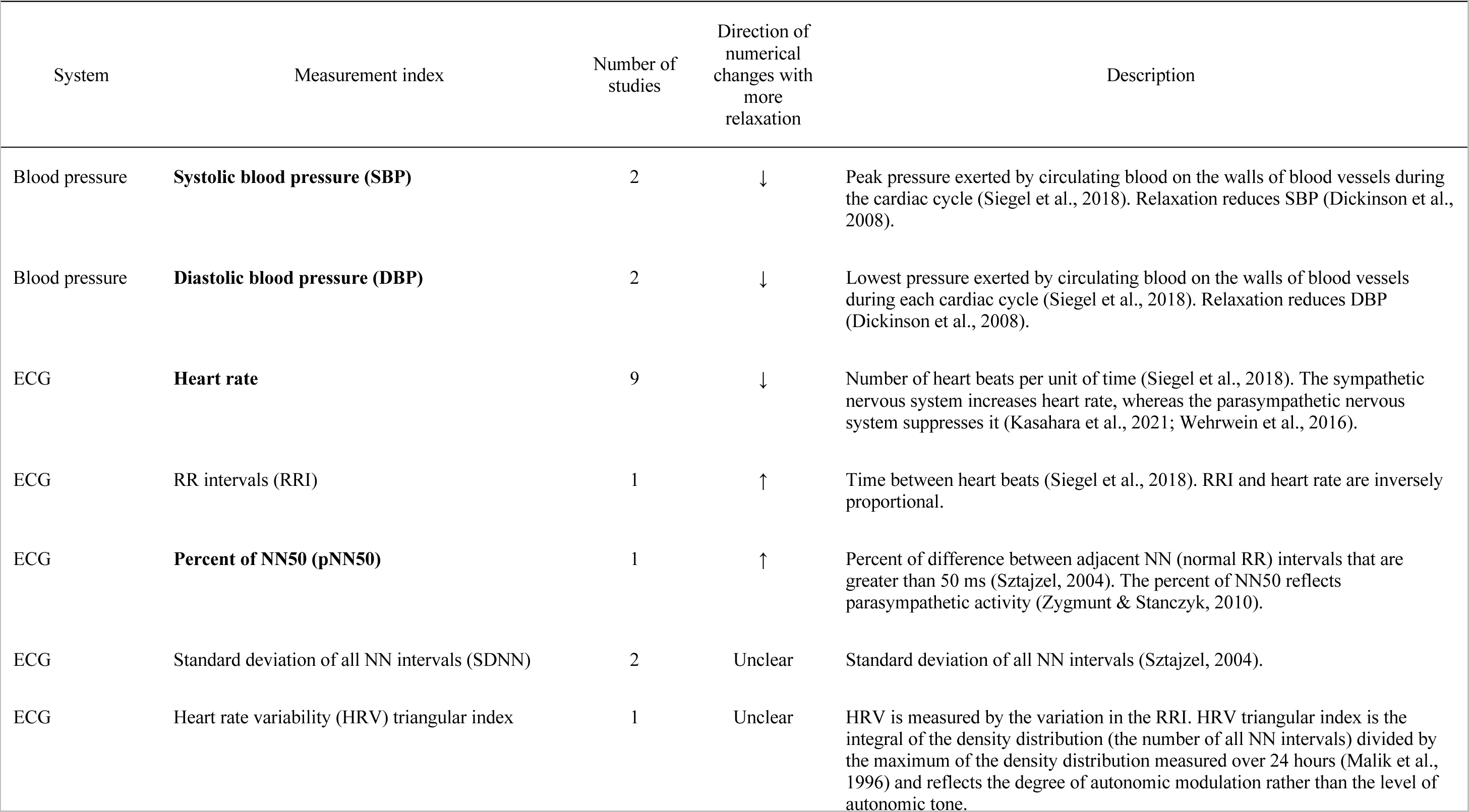
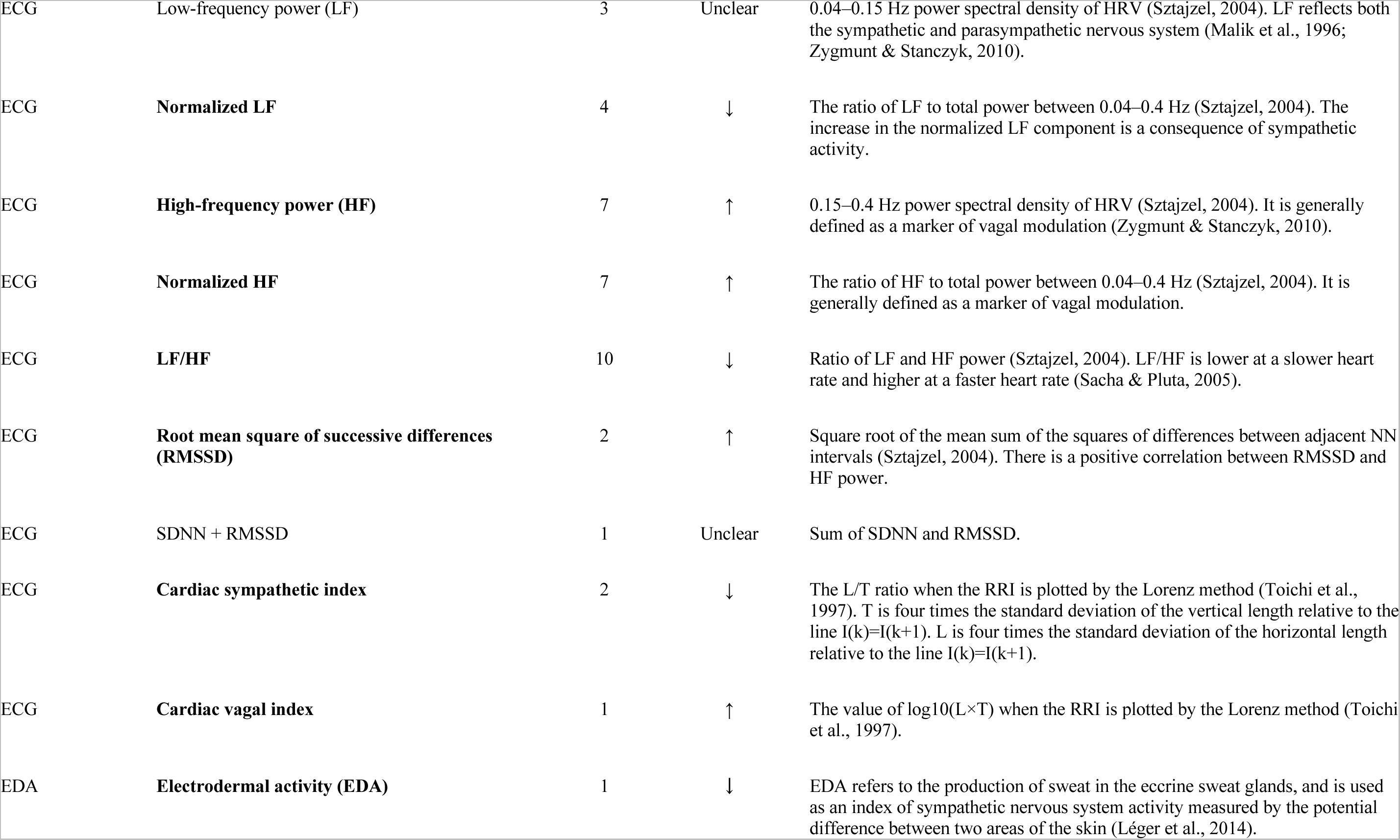
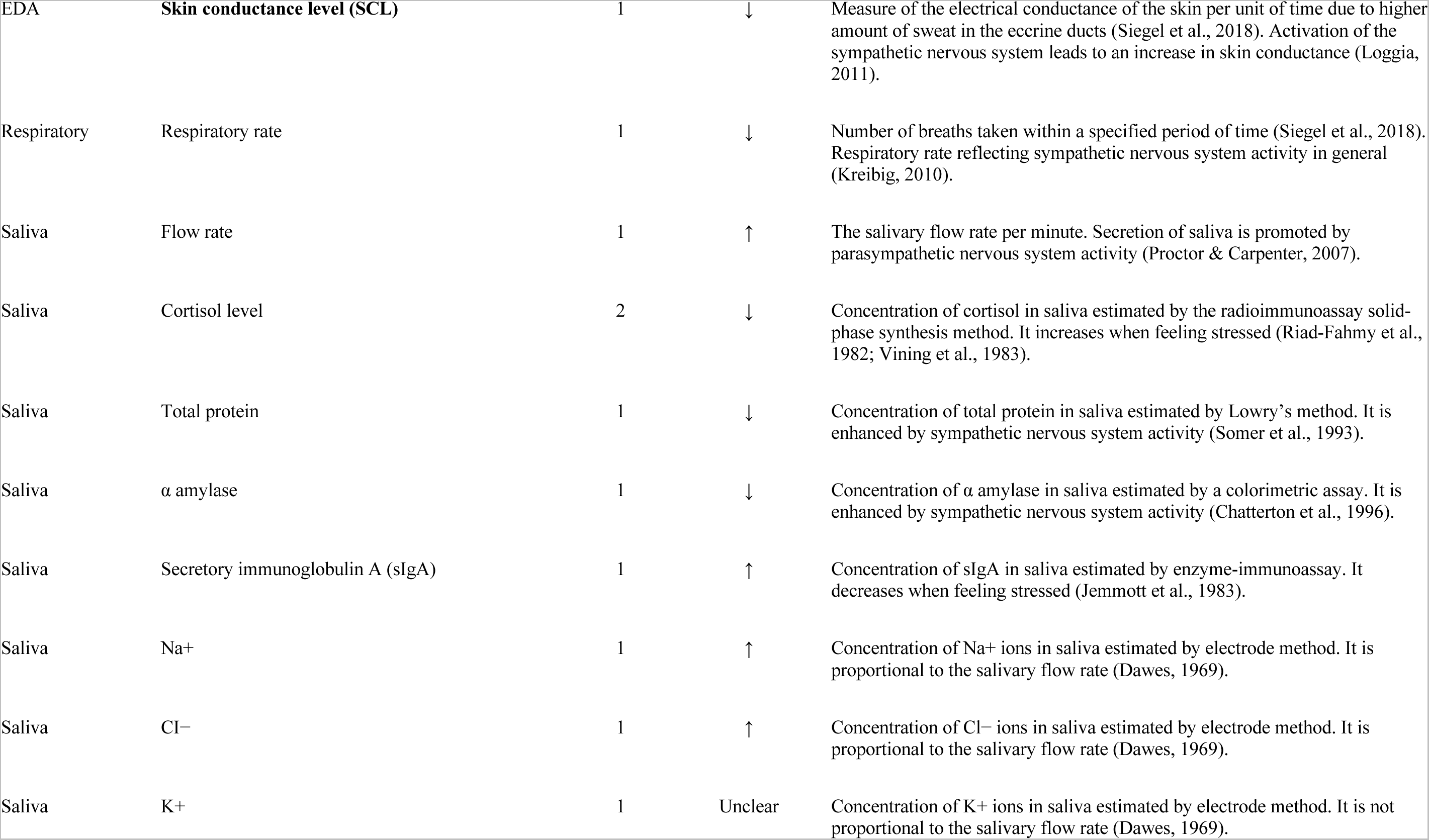
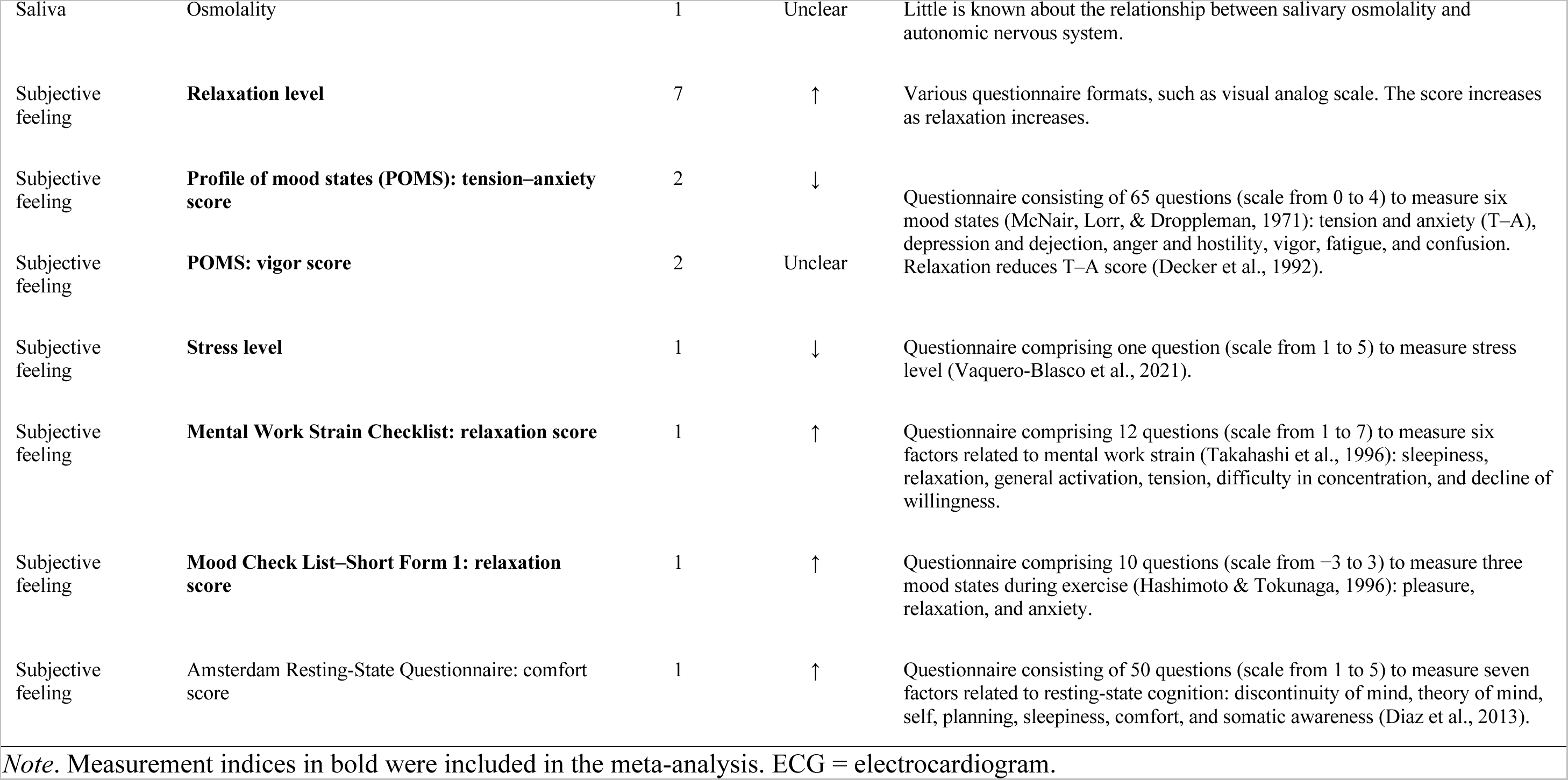
Relaxation indices identified through the qualitative assessment.

#### Effect Size Calculation

To compare effect sizes across studies, Fisher’s transformation was applied to the sign-adjusted *r*-values prior to the meta-analysis. Fisher’s transformation is preferred as the effect size adjustment for meta-analyses with correlation coefficients because it transforms skewed data into an approximately normal distribution. To address the violation of independence assumption, which assumes that effect size estimates from different studies are independent and have sampling distributions with known conditional variances (Hedges et al., 2010; Tipton & Pustejovsky, 2015), the robust variance estimation method was employed to correct Fisher’s transformed *z*-value (Fisher’s *z*) for multiple effect sizes extracted from a single study.

Different frequency bands on the EEG are considered to represent distinct neuronal functions (da Silva, 1991; da Silva, 2013; Sauseng & Klimesch, 2008), thus addressing heterogeneity in frequency band measurements is crucial for accurate effect size estimation. Consequently, the meta-analysis was conducted independently for each frequency band of EEG/MEG indices (delta, theta, alpha, beta, and gamma). Some studies utilized EEG indices of the power ratio between multiple frequency bands. In such instances, we conducted meta-analyses only when a sufficient number of trials and outcome variables were identified.

Fisher’s *z* and its variance were calculated with *r*-value, sample size, and number of reported *r*-values in each study. A standard pairwise meta-analysis was conducted using a random-effects model to combine all *z*-values into a single dataset, assuming heterogeneity between studies, as estimated effect sizes were expected to differ from true effect sizes. Consequently, estimated effect sizes and 95% confidence intervals (CIs) were obtained. If the CI did not include zero, the null hypothesis of no correlation between EEG/MEG and reference indices was rejected.

Heterogeneity was concluded when the Cochrane’s *Q* statistical test yielded a significance level of *p* < .1 (Higgins & Green, 2008). If heterogeneity is observed, the estimated effect sizes do not accurately reflect the true effect sizes due to the lack of appropriate grouping. Effect size and heterogeneity were calculated using the metafor package in R. Robust variance estimation was also conducted using the metafor and clubSandwich packages in R.

#### Subgroup Meta-analysis

When heterogeneity was confirmed, we organized the data into subgroups based on the type of EEG feature components because different feature components may represent distinct biological states. If heterogeneity remained, we classified them into subgroups based on EEG recording channels and type of reference relaxation index. The EEG channel locations were categorized into six regional groups (Sauseng & Klimesch, 2008): prefrontal, frontal, central, parietal, occipital, and temporal. This subdivision is essential for accurate effect size estimation because EEG signals measured from different channels likely reflect the activity of distinct neuronal populations. The type of reference index is also critical for effect size estimation, as measured components may differ between reference indices. For instance, it was reported that introspective components quantified by psychological assessments did not correlate with objective components by physiological assessment (Hites & Lundervold, 2013; Steghaus & Poth, 2022). Furthermore, if heterogeneity persisted within these subgroups, we further divided them into combination subgroups that considered both EEG recording channels and reference indices. Subgroups with an insufficient number of studies (<2) were excluded from the subgroup analysis as meta-analysis was not feasible.

In addition, omnibus tests were performed to verify the null hypothesis of no heterogeneity among the subgroups (Rubio-Aparicio et al., 2017; Rubio-Aparicio et al., 2020). The individual alpha level was adjusted using the Bonferroni method to control the overall experiment-wise error rate.

### Publication Bias Assessment

Publication bias was assessed visually with funnel plots and statistically using Egger’s test (Egger et al., 1997) and Begg’s test (Begg & Mazumdar, 1994). To aggregate effect sizes within the same study, the *escalc*, *vcalc*, *rma.mv*, and *aggregate* functions in metafor were used. The significance for publication bias was set at *p* < .1 for Egger’s and Begg’s tests. Visualization with funnel plots and statistical tests were conducted separately to verify the presence of publication bias within each frequency category.

### Transparency and Openness

We adhered to the PRISMA 2020 guidelines for systematic reviews (Page et al., 2021) and the MARS guidelines for meta-analytic reporting (Appelbaum et al., 2018). All data, analysis codes, and research materials (including our coding scheme) are available at <u>the OSF repository</u>. Data were analyzed using R version 4.2.3 (R Core Team, 2021), the metafor package version 3.0-2 (Viechtbauer, 2010), and the clubSandwich package version 0.5.8. This review project was not preregistered.

## Results

### Search Results

The workflow of the literature review is depicted in Fig. 1. We initially retrieved 3,295 studies from the databases, of which 2,943 were excluded during the first screen based on title and abstract reviews. During the second full-text screen, we excluded an additional 314 studies, including 237 that did not report correlation coefficients, 62 that did not measure relaxation, nine lacking EEG/MEG measurements, three enrolling subjects younger than 18 or older than 65 years, two lacking measurements while awake, and one including patients. As a result, 38 studies with 1,120 total participants were included in the qualitative synthesis. None of these studies used magnetoencephalography (MEG).

**Figure 1.**
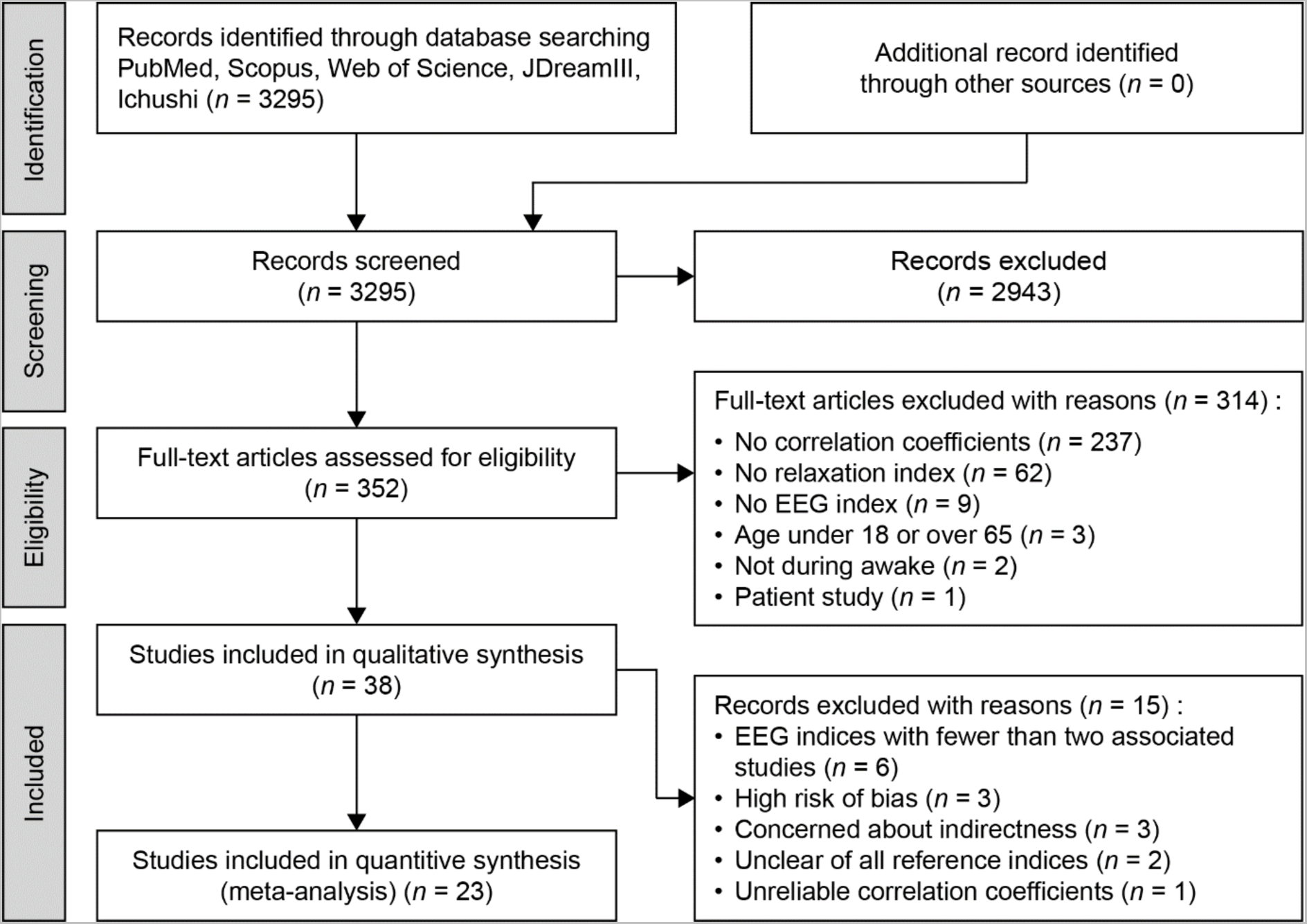
PRISMA flowchart of literature search and inclusion. *n* = the number of studies.

In addition, we excluded nine studies from the meta-analysis for high risk of biases (*n* = 3), concerns about indirectness (*n* = 3), unclear directions of numerical changes with greater relaxation for all reported reference indices (*n* = 2), and unreliable correlation coefficients (*n* = 1). Additionally, six studies that only used indices with less than two studies employing the respective indices (e.g., microstate, beta/alpha ratio) were excluded. Finally, 23 studies were retained for the meta-analysis.

### Study and Sample Characteristics

The main characteristics of the included studies are summarized in Table 2. The publication years spanned from 1976 to 2021, with over half published after 2014 (Fig. 2A). Frontal channels (F3, F4) and occipital channels (O1, O2) were most often used for EEG recording, followed by the left and right central channels (C3, C4), and then the prefrontal (Fp1, Fp2) and parietal channels (P3, P4) (Fig. 2B), while midline and edge channels were rarely used. The alpha band was the most common band of interest, followed by the theta and beta bands (Fig. 2C), while delta (<4 Hz) and gamma bands were less often included in the analysis. Other measures utilizing multiple frequency bands, such as the alpha/beta power ratio, were also infrequently reported. The majority of studies reported band power or normalized power, although various other EEG components were also reported (Fig. 2D).

**Figure 2.**
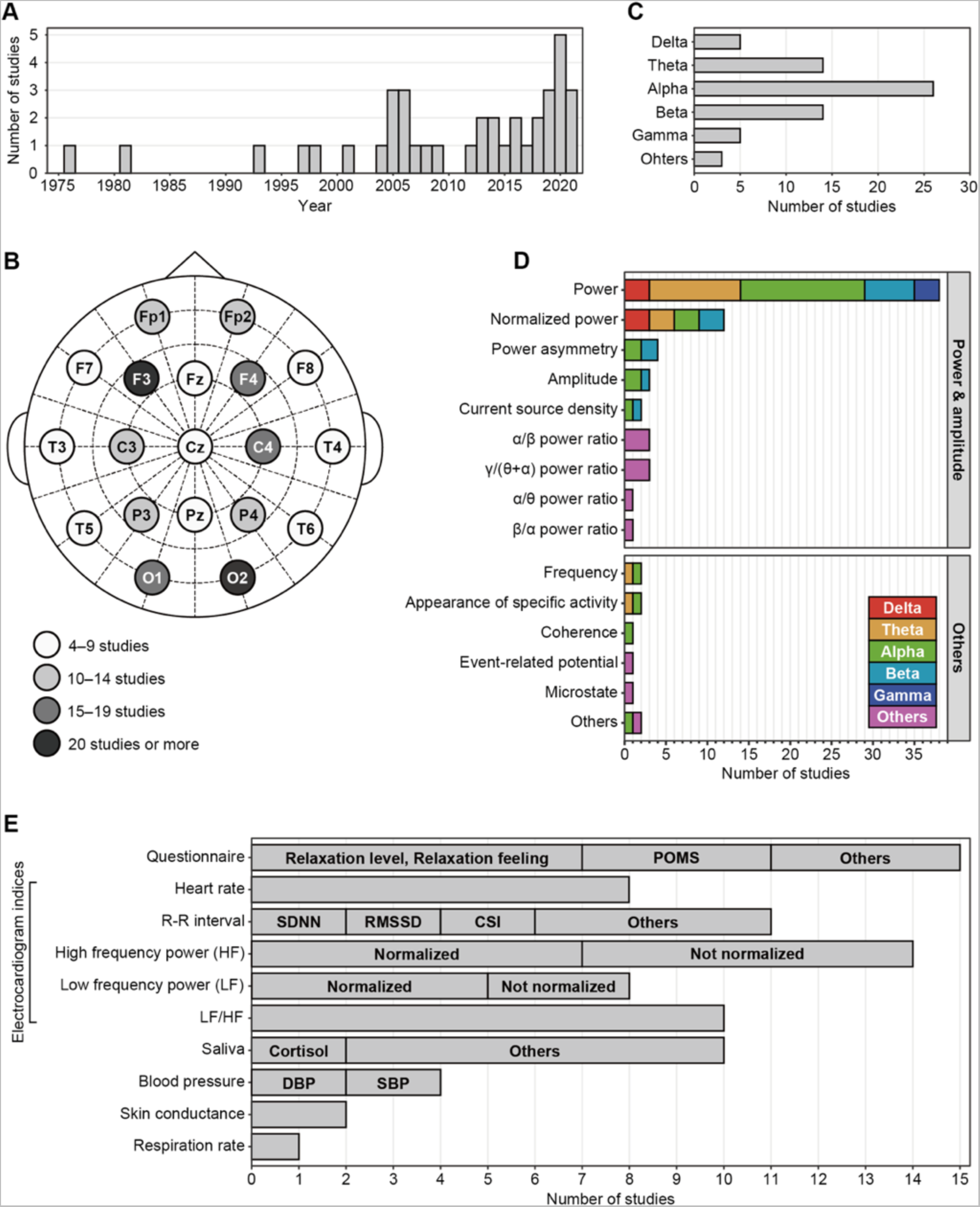
Qualitative assessment of the 38 eligible studies. (A) Number of studies from each publication year. (B) Number of studies using the indicated EEG recording channel locations. (C) Number of studies reporting the power of the indicated EEG frequency band. (D) EEG feature components measured. (E) Reference indices measured. CSI = cardiac sympathetic index; DBP = diastolic blood pressure; POMS = profile of mood states; RMSSD = root mean square of successive differences; SBP = systolic blood pressure; SDNN = standard deviation of NN intervals.

**Table 2.**
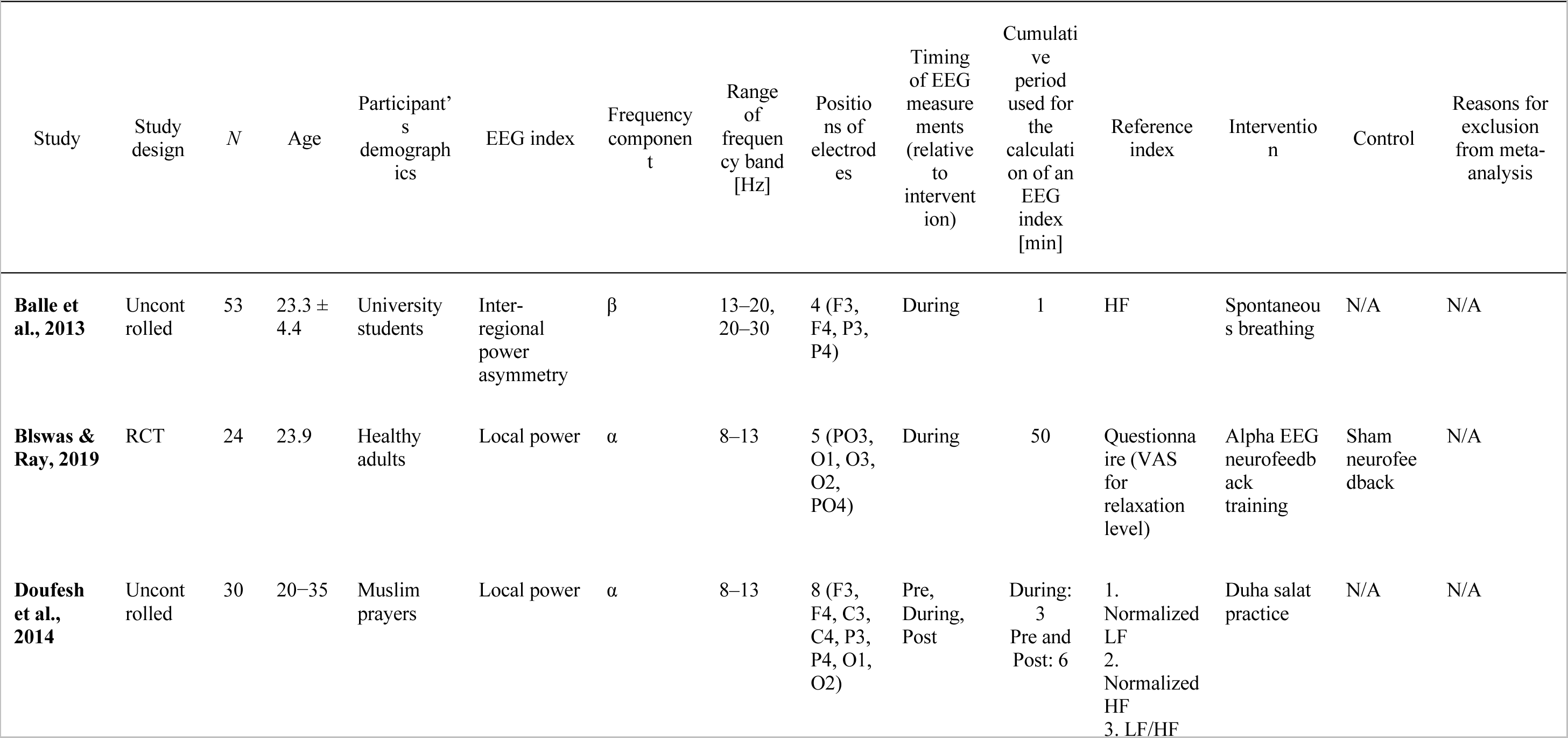
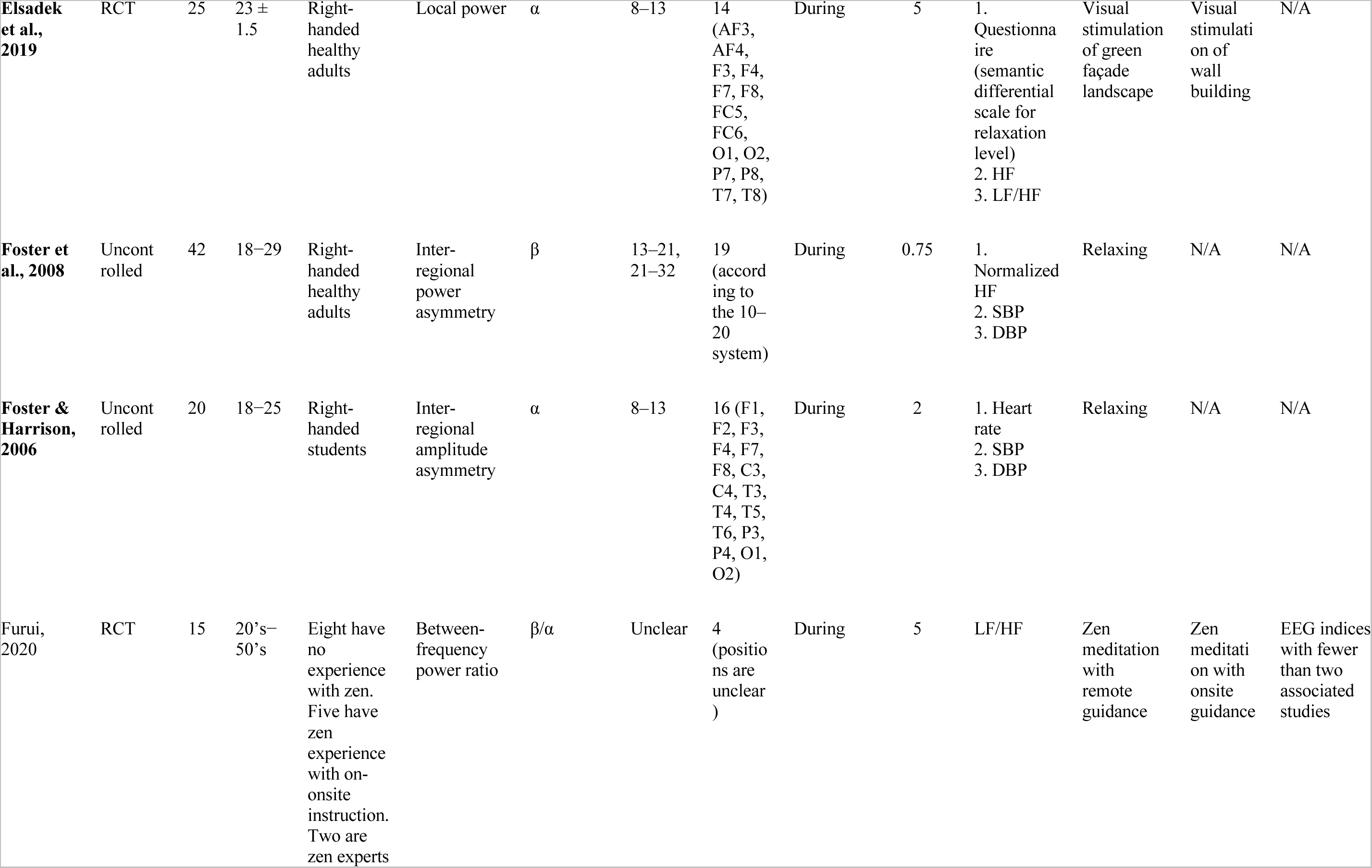
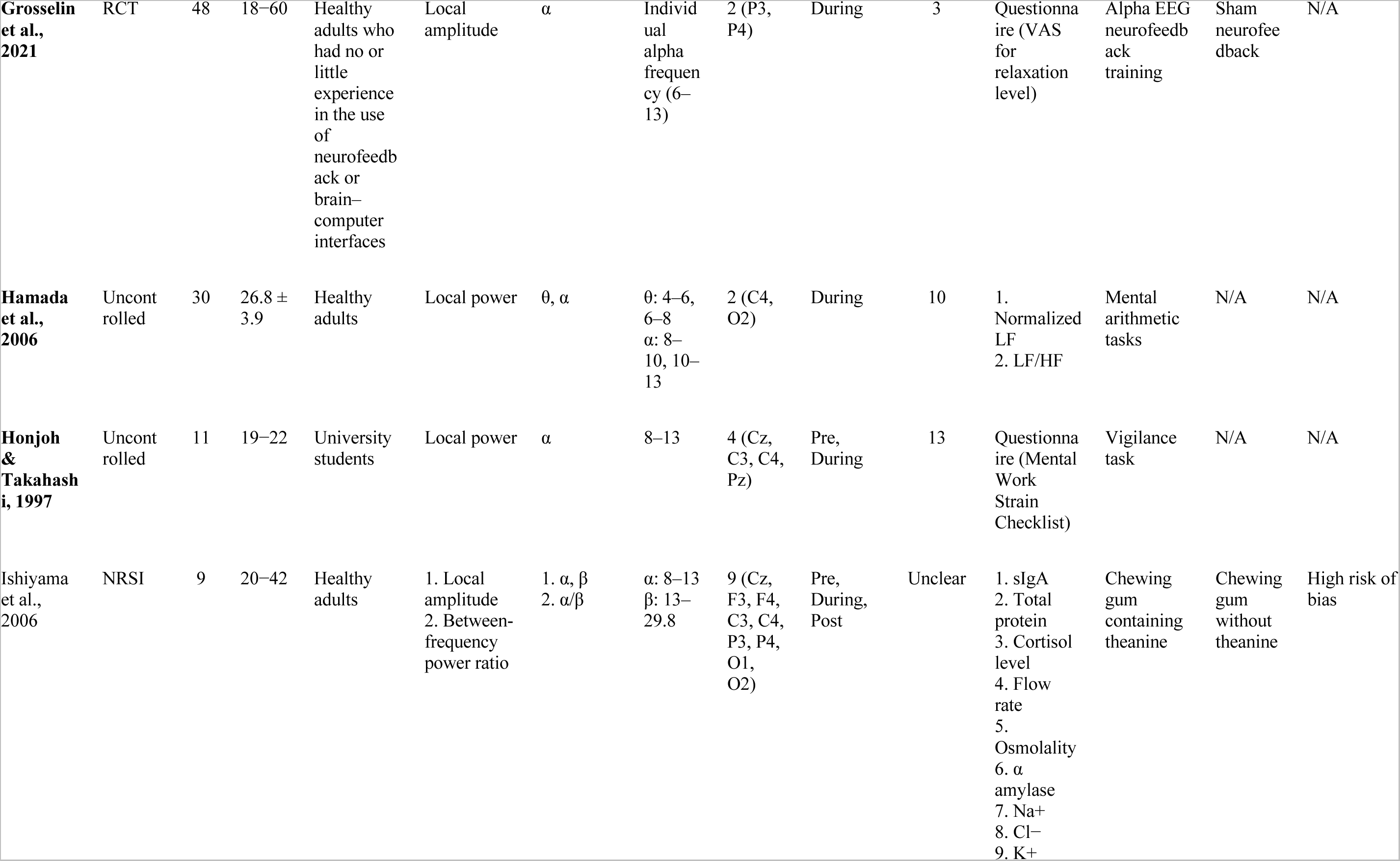
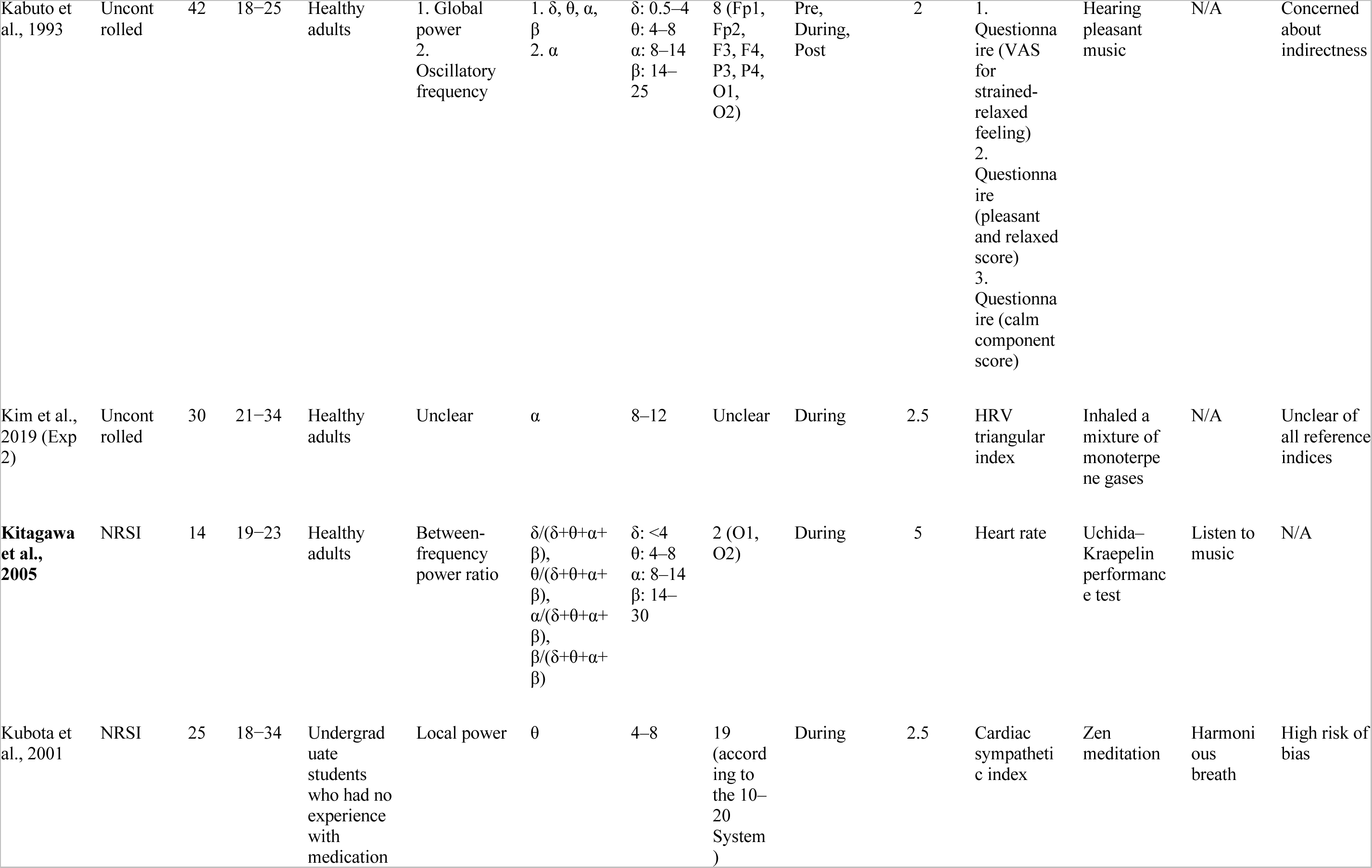
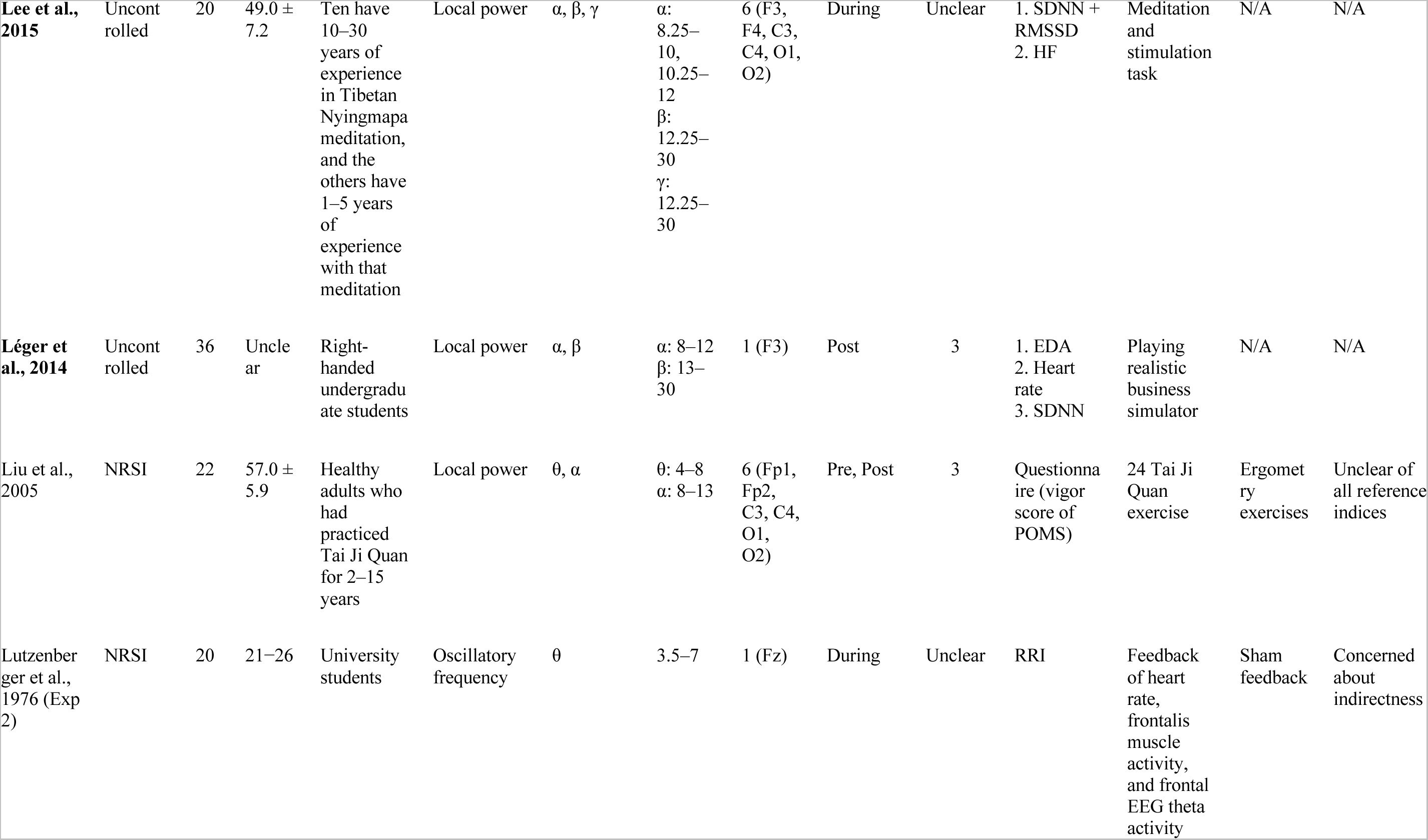
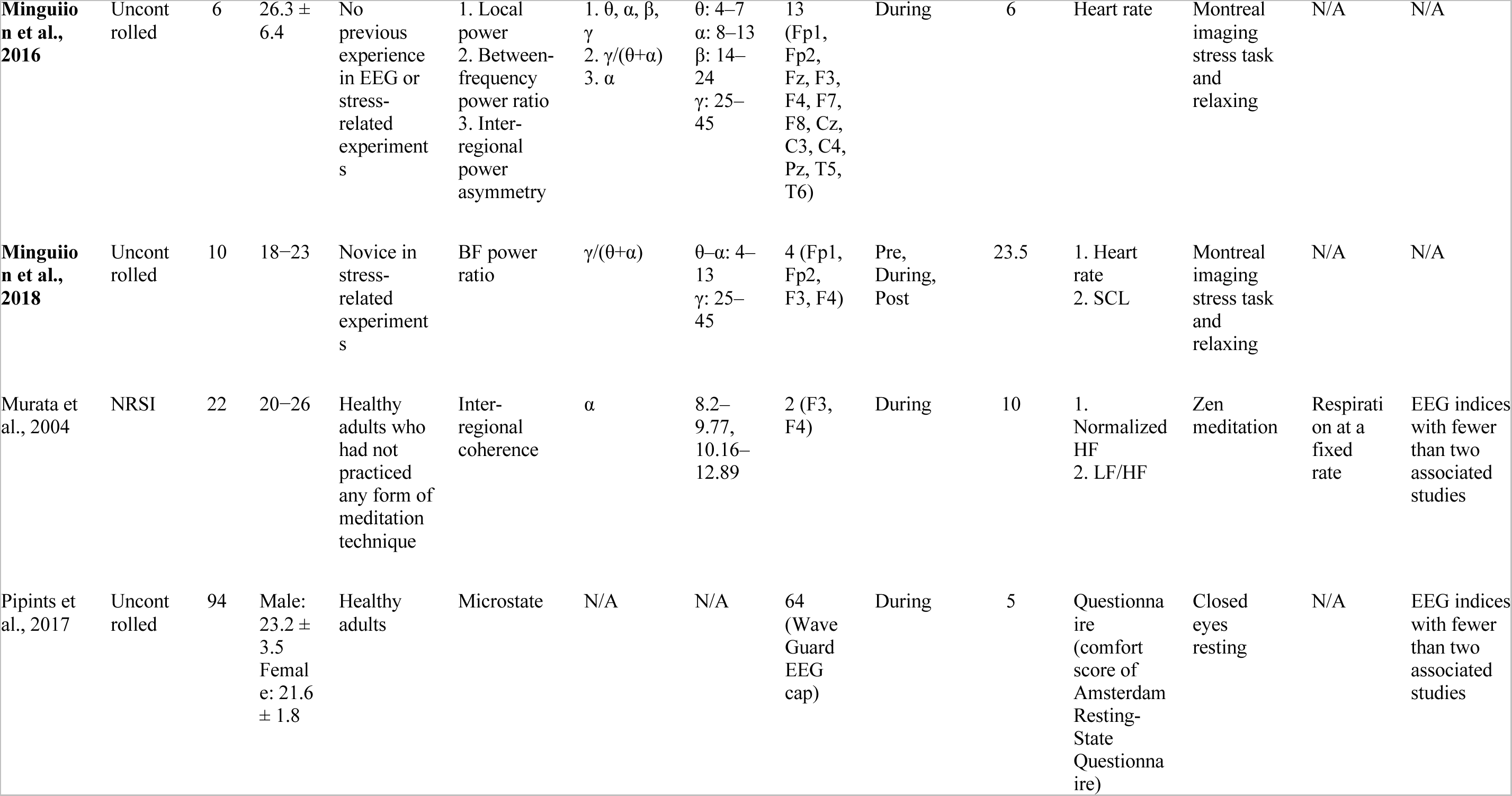
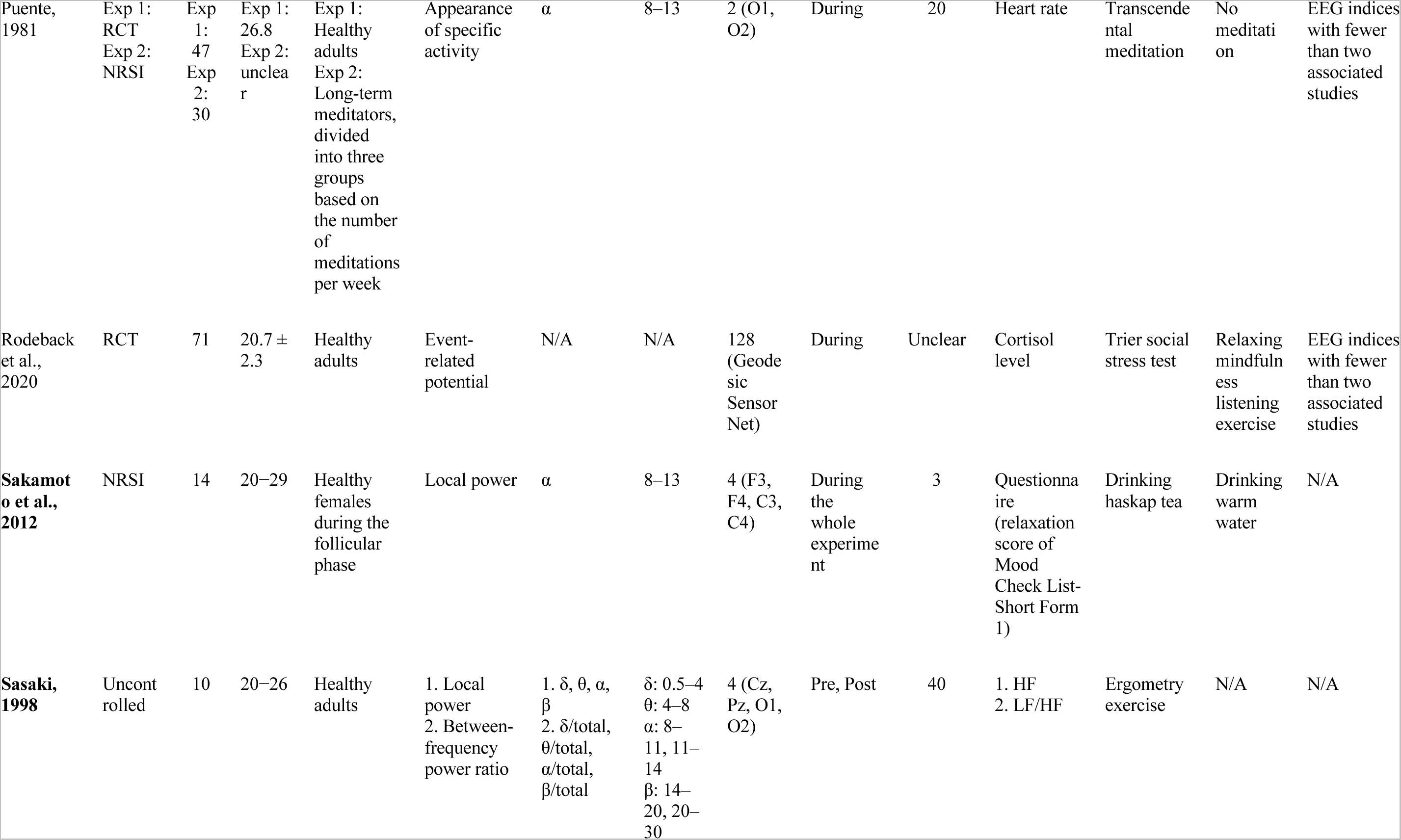
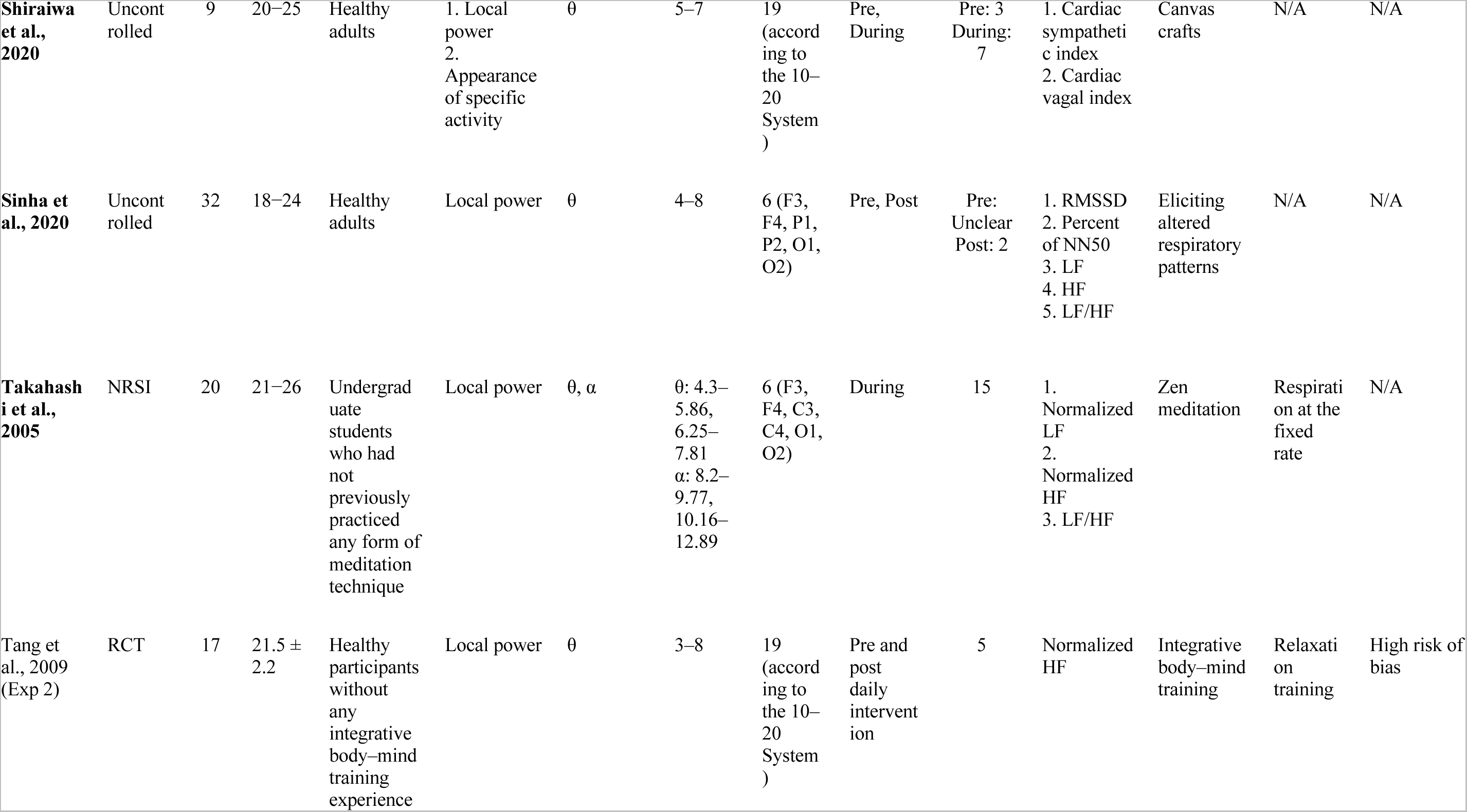
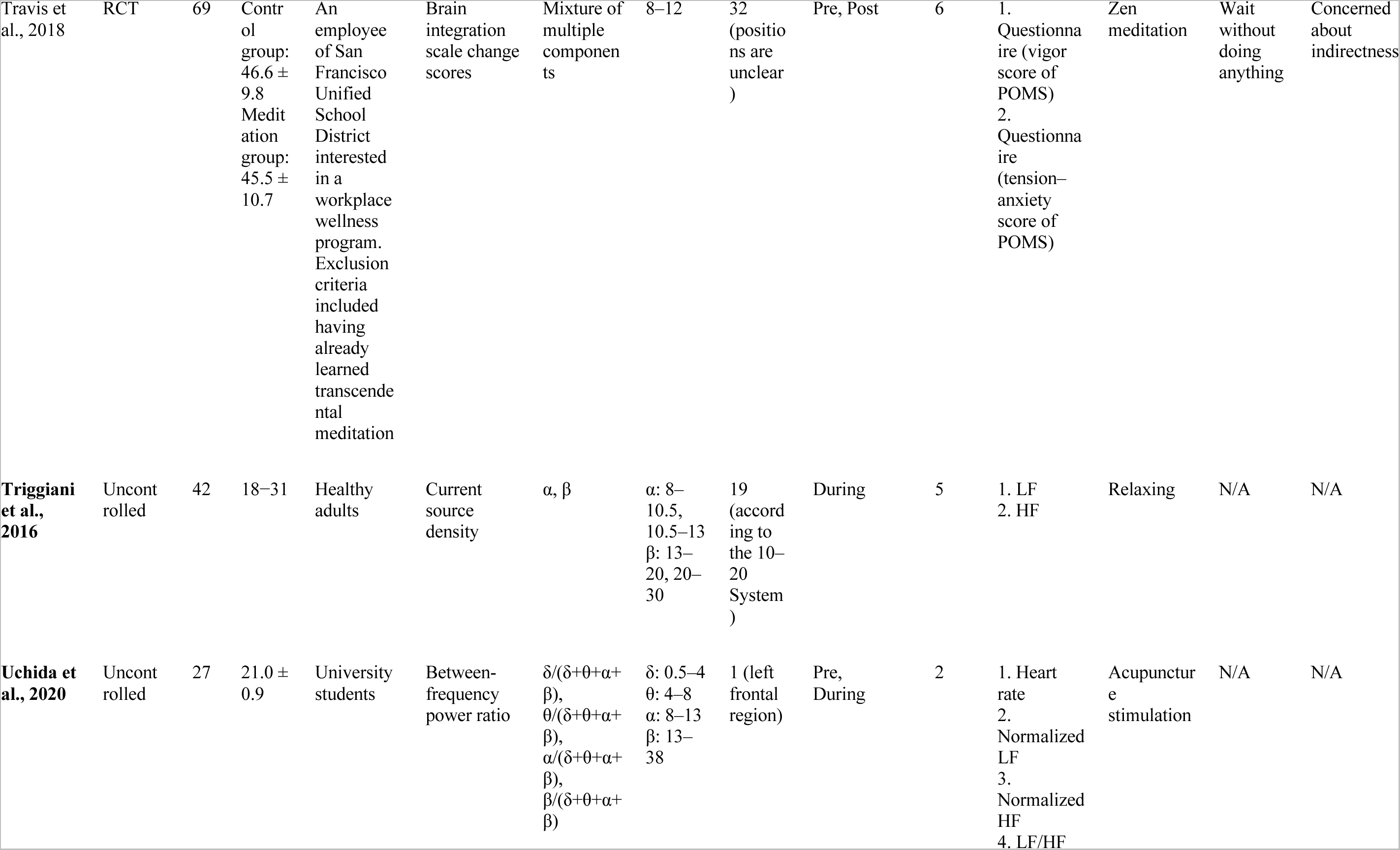
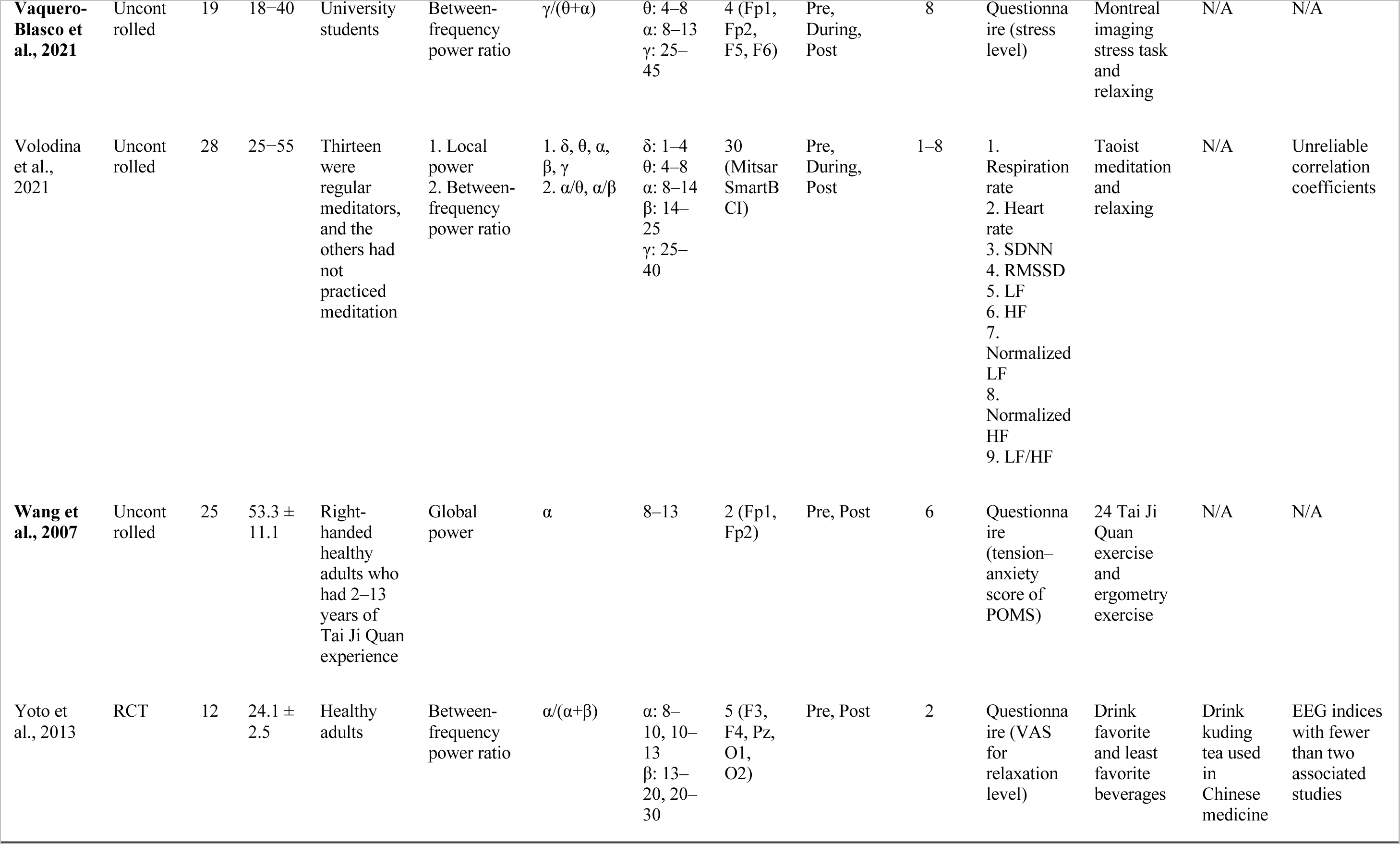
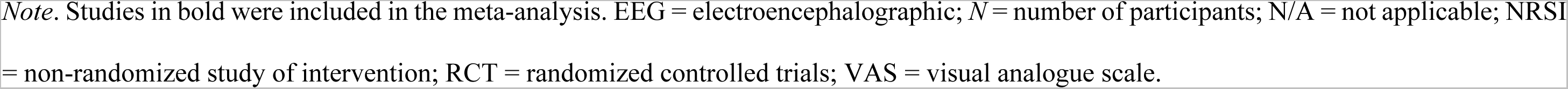
Characteristics of the studies included in the qualitative synthesis.

| Study | Study design | *N* | Age | Participant's demographics | EEG index | Frequency component | Range of frequency band [Hz] | Positions of electrodes | Timing of EEG measurements (relative to intervention) | Cumulative period used for the calculation of an EEG index [min] | Reference index | Intervention | Control | Reasons for exclusion from meta-analysis |
| --- | --- | --- | --- | --- | --- | --- | --- | --- | --- | --- | --- | --- | --- | --- |
| **Balle et al., 2013** | Uncontrolled | 53 | 23.3 ± 4.4 | University students | Inter-regional power asymmetry | $\beta$ | 13–20, 20–30 | 4 (F3, F4, P3, P4) | During | 1 | HF | Spontaneous breathing | N/A | N/A |
| **Blswas & Ray, 2019** | RCT | 24 | 23.9 | Healthy adults | Local power | $\alpha$ | 8–13 | 5 (PO3, O1, O3, O2, PO4) | During | 50 | Questionnaire (VAS for relaxation level) | Alpha EEG neurofeedback training | Sham neurofeedback | N/A |
| **Doufesh et al., 2014** | Uncontrolled | 30 | 20–35 | Muslim prayers | Local power | $\alpha$ | 8–13 | 8 (F3, F4, C3, C4, P3, P4, O1, O2) | Pre, During, Post | During: 3<br>Pre and Post: 6 | 1. Normalized LF<br>2. Normalized HF<br>3. LF/HF | Duha salat practice | N/A | N/A |
| **Elsadek et al., 2019** | RCT | 25 | 23 ± 1.5 | Right-handed healthy adults | Local power | $\alpha$ | 8–13 | 14 (AF3, AF4, F3, F4, F7, F8, FC5, FC6, O1, O2, P7, P8, T7, T8) | During | 5 | 1. Questionnaire (semantic differential scale for relaxation level)<br>2. HF<br>3. LF/HF | Visual stimulation of green façade landscape | Visual stimulation of wall building | N/A |
| **Foster et al., 2008** | Uncontrolled | 42 | 18–29 | Right-handed healthy adults | Inter-regional power asymmetry | $\beta$ | 13–21, 21–32 | 19 (according to the 10–20 system) | During | 0.75 | 1. Normalized HF<br>2. SBP<br>3. DBP | Relaxing | N/A | N/A |
| **Foster & Harrison, 2006** | Uncontrolled | 20 | 18–25 | Right-handed students | Inter-regional amplitude asymmetry | $\alpha$ | 8–13 | 16 (F1, F2, F3, F4, F7, F8, C3, C4, T3, T4, T5, T6, P3, P4, O1, O2) | During | 2 | 1. Heart rate<br>2. SBP<br>3. DBP | Relaxing | N/A | N/A |
| **Furui, 2020** | RCT | 15 | 20's–50's | Eight have no experience with zen. Five have zen experience with on-site instruction. Two are zen experts | Between-frequency power ratio | $\beta/\alpha$ | Unclear | 4 (positions are unclear) | During | 5 | LF/HF | Zen meditation with remote guidance | Zen meditation with onsite guidance | EEG indices with fewer than two associated studies |
| **Grosselin et al., 2021** | RCT | 48 | 18–60 | Healthy adults who had no or little experience in the use of neurofeedback or brain–computer interfaces | Local amplitude | $\alpha$ | Individual alpha frequency (6–13) | 2 (P3, P4) | During | 3 | Questionnaire (VAS for relaxation level) | Alpha EEG neurofeedback training | Sham neurofeedback | N/A |
| **Hamada et al., 2006** | Uncontrolled | 30 | 26.8 ± 3.9 | Healthy adults | Local power | $\theta, \alpha$ | $\theta$: 4–6, 6–8<br>$\alpha$: 8–10, 10–13 | 2 (C4, O2) | During | 10 | 1. Normalized LF<br>2. LF/HF | Mental arithmetic tasks | N/A | N/A |
| **Honjoh & Takahashi, 1997** | Uncontrolled | 11 | 19–22 | University students | Local power | $\alpha$ | 8–13 | 4 (Cz, C3, C4, Pz) | Pre, During | 13 | Questionnaire (Mental Work Strain Checklist) | Vigilance task | N/A | N/A |
| Ishiyama et al., 2006 | NRSI | 9 | 20–42 | Healthy adults | 1. Local amplitude<br>2. Between-frequency power ratio | 1. $\alpha, \beta$<br>2. $\alpha/\beta$ | $\alpha$: 8–13<br>$\beta$: 13–29.8 | 9 (Cz, F3, F4, C3, C4, P3, P4, O1, O2) | Pre, During, Post | Unclear | 1. sIgA<br>2. Total protein<br>3. Cortisol level<br>4. Flow rate<br>5. Osmolality<br>6. $\alpha$ amylase<br>7. Na+<br>8. Cl−<br>9. K+ | Chewing gum containing theanine | Chewing gum without theanine | High risk of bias |
| Kabuto et al., 1993 | Uncontrolled | 42 | 18–25 | Healthy adults | 1. Global power<br>2. Oscillatory frequency | 1. $\delta$, $\theta$, $\alpha$, $\beta$<br>2. $\alpha$ | $\delta$: 0.5–4<br>$\theta$: 4–8<br>$\alpha$: 8–14<br>$\beta$: 14–25 | 8 (Fp1, Fp2, F3, F4, P3, P4, O1, O2) | Pre, During, Post | 2 | 1. Questionnaire (VAS for strained-relaxed feeling)<br>2. Questionnaire (pleasant and relaxed score)<br>3. Questionnaire (calm component score) | Hearing pleasant music | N/A | Concerned about indirectness |
| Kim et al., 2019 (Exp 2) | Uncontrolled | 30 | 21–34 | Healthy adults | Unclear | $\alpha$ | 8–12 | Unclear | During | 2.5 | HRV triangular index | Inhaled a mixture of monoterpenic gases | N/A | Unclear of all reference indices |
| Kitagawa et al., 2005 | NRSI | 14 | 19–23 | Healthy adults | Between-frequency power ratio | $\delta/(\delta+\theta+\alpha+\beta)$,<br>$\theta/(\delta+\theta+\alpha+\beta)$,<br>$\alpha/(\delta+\theta+\alpha+\beta)$,<br>$\beta/(\delta+\theta+\alpha+\beta)$ | $\delta$: <4<br>$\theta$: 4–8<br>$\alpha$: 8–14<br>$\beta$: 14–30 | 2 (O1, O2) | During | 5 | Heart rate | Uchida–Kraepelin performance test | Listen to music | N/A |
| Kubota et al., 2001 | NRSI | 25 | 18–34 | Undergraduate students who had no experience with medication | Local power | $\theta$ | 4–8 | 19 (according to the 10–20 System) | During | 2.5 | Cardiac sympathetic index | Zen meditation | Harmonious breath | High risk of bias |
| **Lee et al., 2015** | Uncontrolled | 20 | 49.0 ± 7.2 | Ten have 10–30 years of experience in Tibetan Nyingmapa meditation, and the others have 1–5 years of experience with that meditation | Local power | $\alpha, \beta, \gamma$ | $\alpha$: 8.25–10, 10.25–12<br>$\beta$: 12.25–30<br>$\gamma$: 12.25–30 | 6 (F3, F4, C3, C4, O1, O2) | During | Unclear | 1. SDNN + RMSSD<br>2. HF | Meditation and stimulation task | N/A | N/A |
| **Léger et al., 2014** | Uncontrolled | 36 | Unclear | Right-handed undergraduate students | Local power | $\alpha, \beta$ | $\alpha$: 8–12<br>$\beta$: 13–30 | 1 (F3) | Post | 3 | 1. EDA<br>2. Heart rate<br>3. SDNN | Playing realistic business simulator | N/A | N/A |
| Liu et al., 2005 | NRSI | 22 | 57.0 ± 5.9 | Healthy adults who had practiced Tai Ji Quan for 2–15 years | Local power | $\theta, \alpha$ | $\theta$: 4–8<br>$\alpha$: 8–13 | 6 (Fp1, Fp2, C3, C4, O1, O2) | Pre, Post | 3 | Questionnaire (vigor score of POMS) | 24 Tai Ji Quan exercise | Ergometry exercises | Unclear of all reference indices |
| Lutzenberger et al., 1976 (Exp 2) | NRSI | 20 | 21–26 | University students | Oscillatory frequency | $\theta$ | 3.5–7 | 1 (Fz) | During | Unclear | RRI | Feedback of heart rate, frontalis muscle activity, and frontal EEG theta activity | Sham feedback | Concerned about indirectness |
| **Minguiio**<br>**n et al.,**<br>**2016** | Uncont<br>rolled | 6 | 26.3 ±<br>6.4 | No<br>previous<br>experience<br>in EEG or<br>stress-<br>related<br>experiment<br>s | 1. Local<br>power<br>2. Between-<br>frequency<br>power ratio<br>3. Inter-<br>regional<br>power<br>asymmetry | 1. $\theta$, $\alpha$, $\beta$,<br>$\gamma$<br>2. $\gamma/(\theta+\alpha)$<br>3. $\alpha$ | $\theta$: 4–7<br>$\alpha$: 8–13<br>$\beta$: 14–<br>24<br>$\gamma$: 25–<br>45 | 13<br>(Fp1,<br>Fp2,<br>Fz, F3,<br>F4, F7,<br>F8, Cz,<br>C3, C4,<br>Pz, T5,<br>T6) | During | 6 | Heart rate | Montreal<br>imaging<br>stress task<br>and<br>relaxing | N/A | N/A |
| **Minguiio**<br>**n et al.,**<br>**2018** | Uncont<br>rolled | 10 | 18–23 | Novice in<br>stress-<br>related<br>experiment<br>s | BF power<br>ratio | $\gamma/(\theta+\alpha)$ | θ–α: 4–<br>13<br>$\gamma$: 25–<br>45 | 4 (Fp1,<br>Fp2,<br>F3, F4) | Pre,<br>During,<br>Post | 23.5 | 1. Heart<br>rate<br>2. SCL | Montreal<br>imaging<br>stress task<br>and<br>relaxing | N/A | N/A |
| Murata et<br>al., 2004 | NRSI | 22 | 20–26 | Healthy<br>adults who<br>had not<br>practiced<br>any form of<br>meditation<br>technique | Inter-regional<br>coherence | $\alpha$ | 8.2–<br>9.77,<br>10.16–<br>12.89 | 2 (F3,<br>F4) | During | 10 | 1. Normalized<br>HF<br>2. LF/HF | Zen<br>meditation | Respirati<br>on at a<br>fixed<br>rate | EEG indices<br>with fewer<br>than two<br>associated<br>studies |
| Pipints et<br>al., 2017 | Uncont<br>rolled | 94 | Male:<br>23.2 ±<br>3.5<br>Femal<br>e: 21.6<br>± 1.8 | Healthy<br>adults | Microstate | N/A | N/A | 64<br>(Wave<br>Guard<br>EEG<br>cap) | During | 5 | Questionna<br>ire<br>(comfort<br>score of<br>Amsterdam<br>Resting-State<br>Questionna<br>ire) | Closed<br>eyes<br>resting | N/A | EEG indices<br>with fewer<br>than two<br>associated<br>studies |
| Puente, 1981 | Exp 1: RCT<br>Exp 2: NRSI | Exp 1: 47<br>Exp 2: 30 | Exp 1: 26.8<br>Exp 2: unclear | Exp 1: Healthy adults<br>Exp 2: Long-term meditators, divided into three groups based on the number of meditations per week | Appearance of specific activity | $\alpha$ | 8–13 | 2 (O1, O2) | During | 20 | Heart rate | Transcendental meditation | No meditation | EEG indices with fewer than two associated studies |
| Rodeback et al., 2020 | RCT | 71 | 20.7 ± 2.3 | Healthy adults | Event-related potential | N/A | N/A | 128 (Geodesic Sensor Net) | During | Unclear | Cortisol level | Trier social stress test | Relaxing mindfulness listening exercise | EEG indices with fewer than two associated studies |
| **Sakamoto et al., 2012** | NRSI | 14 | 20–29 | Healthy females during the follicular phase | Local power | $\alpha$ | 8–13 | 4 (F3, F4, C3, C4) | During the whole experiment | 3 | Questionnaire (relaxation score of Mood Check List-Short Form 1) | Drinking haskap tea | Drinking warm water | N/A |
| **Sasaki, 1998** | Uncontrolled | 10 | 20–26 | Healthy adults | 1. Local power<br>2. Between-frequency power ratio | 1. $\delta$, $\theta$, $\alpha$, $\beta$<br>2. $\delta$/total, $\theta$/total, $\alpha$/total, $\beta$/total | $\delta$: 0.5–4<br>$\theta$: 4–8<br>$\alpha$: 8–11, 11–14<br>$\beta$: 14–20, 20–30 | 4 (Cz, Pz, O1, O2) | Pre, Post | 40 | 1. HF<br>2. LF/HF | Ergometry exercise | N/A | N/A |
| **Shiraiwa et al., 2020** | Uncontrolled | 9 | 20–25 | Healthy adults | 1. Local power<br>2. Appearance of specific activity | $\theta$ | 5–7 | 19 (according to the 10–20 System) | Pre, During | Pre: 3<br>During: 7 | 1. Cardiac sympathetic index<br>2. Cardiac vagal index | Canvas crafts | N/A | N/A |
| **Sinha et al., 2020** | Uncontrolled | 32 | 18–24 | Healthy adults | Local power | $\theta$ | 4–8 | 6 (F3, F4, P1, P2, O1, O2) | Pre, Post | Pre: Unclear<br>Post: 2 | 1. RMSSD<br>2. Percent of NN50<br>3. LF<br>4. HF<br>5. LF/HF | Eliciting altered respiratory patterns | N/A | N/A |
| **Takahashi et al., 2005** | NRSI | 20 | 21–26 | Undergraduate students who had not previously practiced any form of meditation technique | Local power | $\theta, \alpha$ | $\theta$: 4.3–5.86,<br>6.25–7.81<br>$\alpha$: 8.2–9.77,<br>10.16–12.89 | 6 (F3, F4, C3, C4, O1, O2) | During | 15 | 1. Normalized LF<br>2. Normalized HF<br>3. LF/HF | Zen meditation | Respiration at the fixed rate | N/A |
| Tang et al., 2009 (Exp 2) | RCT | 17 | 21.5 ± 2.2 | Healthy participants without any integrative body–mind training experience | Local power | $\theta$ | 3–8 | 19 (according to the 10–20 System) | Pre and post daily intervention | 5 | Normalized HF | Integrative body–mind training | Relaxation training | High risk of bias |
| Travis et al., 2018 | RCT | 69 | Control group: 46.6 ± 9.8<br>Meditation group: 45.5 ± 10.7 | An employee of San Francisco Unified School District interested in a workplace wellness program. Exclusion criteria included having already learned transcendental meditation | Brain integration scale change scores | Mixture of multiple components | 8–12 | 32 (positions are unclear) | Pre, Post | 6 | 1. Questionnaire (vigor score of POMS)<br>2. Questionnaire (tension–anxiety score of POMS) | Zen meditation | Wait without doing anything | Concerned about indirectness |
| Triggiani et al., 2016 | Uncontrolled | 42 | 18–31 | Healthy adults | Current source density | $\alpha, \beta$ | $\alpha$: 8–10.5,<br>10.5–13<br>$\beta$: 13–20, 20–30 | 19 (according to the 10–20 System) | During | 5 | 1. LF<br>2. HF | Relaxing | N/A | N/A |
| Uchida et al., 2020 | Uncontrolled | 27 | 21.0 ± 0.9 | University students | Between-frequency power ratio | $\delta/(\delta+\theta+\alpha+\beta)$,<br>$\theta/(\delta+\theta+\alpha+\beta)$,<br>$\alpha/(\delta+\theta+\alpha+\beta)$,<br>$\beta/(\delta+\theta+\alpha+\beta)$ | $\delta$: 0.5–4<br>$\theta$: 4–8<br>$\alpha$: 8–13<br>$\beta$: 13–38 | 1 (left frontal region) | Pre, During | 2 | 1. Heart rate<br>2. Normalized LF<br>3. Normalized HF<br>4. LF/HF | Acupuncture stimulation | N/A | N/A |
| **Vaquero-Blasco et al., 2021** | Uncontrolled | 19 | 18–40 | University students | Between-frequency power ratio | $\gamma/(\theta+\alpha)$ | $\theta$: 4–8<br>$\alpha$: 8–13<br>$\gamma$: 25–45 | 4 (Fp1, Fp2, F5, F6) | Pre, During, Post | 8 | Questionnaire (stress level) | Montreal imaging stress task and relaxing | N/A | N/A |
| Volodina et al., 2021 | Uncontrolled | 28 | 25–55 | Thirteen were regular meditators, and the others had not practiced meditation | 1. Local power<br>2. Between-frequency power ratio | 1. $\delta$, $\theta$, $\alpha$, $\beta$, $\gamma$<br>2. $\alpha/\theta$, $\alpha/\beta$ | $\delta$: 1–4<br>$\theta$: 4–8<br>$\alpha$: 8–14<br>$\beta$: 14–25<br>$\gamma$: 25–40 | 30 (Mitsar SmartB CI) | Pre, During, Post | 1–8 | 1. Respiration rate<br>2. Heart rate<br>3. SDNN<br>4. RMSSD<br>5. LF<br>6. HF<br>7. Normalized LF<br>8. Normalized HF<br>9. LF/HF | Taoist meditation and relaxing | N/A | Unreliable correlation coefficients |
| **Wang et al., 2007** | Uncontrolled | 25 | 53.3 ± 11.1 | Right-handed healthy adults who had 2–13 years of Tai Ji Quan experience | Global power | $\alpha$ | 8–13 | 2 (Fp1, Fp2) | Pre, Post | 6 | Questionnaire (tension–anxiety score of POMS) | 24 Tai Ji Quan exercise and ergometry exercise | N/A | N/A |
| Yoto et al., 2013 | RCT | 12 | 24.1 ± 2.5 | Healthy adults | Between-frequency power ratio | $\alpha/(\alpha+\beta)$ | $\alpha$: 8–10, 10–13<br>$\beta$: 13–20, 20–30 | 5 (F3, F4, Pz, O1, O2) | Pre, Post | 2 | Questionnaire (VAS for relaxation level) | Drink favorite and least favorite beverages | Drink kuding tea used in Chinese medicine | EEG indices with fewer than two associated studies |

*Note.* Studies in bold were included in the meta-analysis. EEG = electroencephalographic; *N* = number of participants; N/A = not applicable; NRSI = non-randomized study of intervention; RCT = randomized controlled trials; VAS = visual analogue scale.

The reference indices included both objective and introspective measures. Objective indices included parameters related to the cardiovascular system, saliva, electrodermal response, and respiration (Table 1), while the introspective indices relied exclusively on questionnaires. The most common ECG-related measures were heart rate, low to high-frequency power ratio (LF/HF), and normalized HF. The most frequently measured introspective index (questionnaire item) was relaxation level, although the exact format varied. Neither saliva contents nor electrodermal response features were reported in more than two studies.

We adjusted the signs of reference indices used in the meta-analysis to indicate an increase with greater relaxation. Components considered to increase with relaxation, such as normalized HF, HF, and root mean square of successive differences between normal heartbeats for cardiovascular system indices and relaxation level from questionnaires are indicated by ↑ in Table 1. Conversely, components considered to decrease with relaxation such as LF/HF, heart rate, and normalized LF for cardiovascular system indices as well as tension–anxiety score of Profile of Mood States and stress level from questionnaires are indicated by ↓ in Table 1. The direction of the change in value with increasing relaxation was also specified for saliva, electrodermal, and respiration indices, although these were not used in the meta-analysis. Other components were categorized as unclear.

### Risk of Bias Assessment

The risk of bias for the 38 eligible studies is visualized in Fig. 3. All studies except for two were determined to have low risk of bias in participant selection. In one of the two studies judged to have unclear risk, information on the control of participant characteristics when assigning participants to experimental groups was not provided. In the other study, participants’ experiences with zen varied and were not controlled for across groups. Twelve articles were rated as unclear regarding risk of bias from confounding variables due to insufficient participant information, missing randomization details, or suspected order effects. Two articles with apparent order effects were rated as high risk. Eight articles lacking descriptions of blinding in measurements or consistency of measurement indices were evaluated as unclear for risk of bias in intervention measurements. Other articles were rated as low risk since the concurrency between EEG and reference index measurements could be confirmed. Four studies in which two authors (HK and KS) calculated the correlation coefficient from the raw data or other statistics were appraised as low risk of bias from outcome assessment. Studies without clear explanations of researcher blinding during outcome assessment were graded as unclear. Six studies lacking descriptions of the analyzed data sample size or not specifying whether all data were in fact analyzed were regarded as unclear for risk of bias due to incomplete outcome data. Nine studies in which the number of excluded participants exceeded 10% in a single group were classified as high risk. Twenty-seven studies were deemed unclear regarding risk of bias from selective reporting because pre-registration was not conducted. Seven studies in which reporting bias was identified, such as reporting only significant results, were categorized as high risk. Other studies for which we calculated correlation coefficients from the raw data or other statistics were deemed low risk. Finally, 34 studies lacking descriptions of sample size setting or COI were ranked as unclear for risk of other sources of bias. Two studies considered to have a potential COI were rated as high risk. Ultimately, three studies judged to have a high risk of bias in more than one domain were excluded from the meta-analysis.

**Figure 3.**
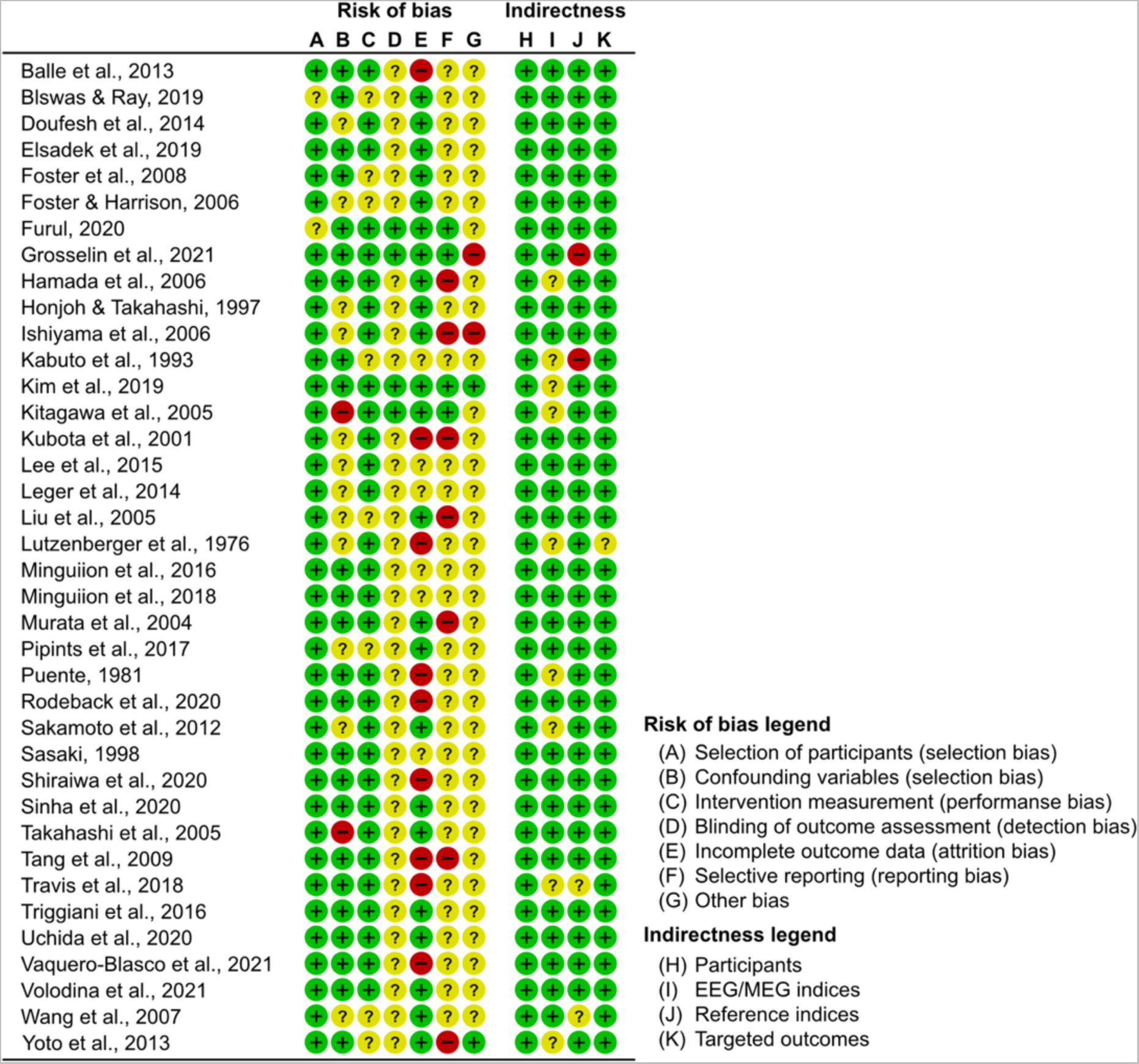
Assessments of risk of bias and indirectness. Green, low risk of bias or low indirectness; red, high risk of bias or high indirectness; yellow, unclear risk of bias or unclear indirectness.

### Indirectness Assessment

Indirectness of the 38 eligible studies is visualized in Fig. 3. For participant selection, all studies were rated as low because they properly described information about participants. Regarding EEG recording and calculation procedures of EEG indices, 23 studies were assessed as low, and nine studies as unclear. For the targeted outcome, all but one studies were rated as low. For the physiological state captured by the EEG index and the relaxed state measured by reference indices, 34 studies were classified as low, two as unclear, and two as high. Only studies that did not provide clear information on the measures used to calculate the correlation coefficients were rated as unclear. Ultimately, three studies judged to have unclear or high indirectness in more than one domain were excluded from the meta-analysis.

### Correlation Between EEG and Reference Relaxation Indices

The meta-analysis exclusively considered EEG indices related to oscillation band power or amplitude (i.e., those listed in the “Power/Amplitude” category in Fig. 2D). Overall, no significant correlations were found between reference indices and any EEG band power or amplitude (Table 3). Notably, there was no heterogeneity in the delta, gamma, and relative gamma bands, indicating that the estimated effect sizes in these categories accurately reflected the real effect sizes. Conversely, significant heterogeneity was found in the theta, alpha, and beta bands. This prompted us to conduct subgroup analyses investigating whether the relationships between EEG and relaxation indices vary depending on the recording channel location and type of reference relaxation index.

**Table 3.** Summary of meta-analysis results on correlations between EEG and reference relaxation indices.

| EEG frequency bands | Number of studies | Number of subjects | Number of outcome measures | Fisher's *z*-transformed correlation coefficient | 95% CI | *Q* |
| --- | --- | --- | --- | --- | --- | --- |
| Delta | 3 | 51 | 18 | −0.04 | [−0.13, 0.05] | 7.01 |
| Theta | 8 | 148 | 46 | 0.22 | [−0.17, 0.62] | 92.72* |
| Alpha | 17 | 402 | 203 | 0.11 | [−0.14, 0.36] | 373.77* |
| Beta | 9 | 250 | 273 | −0.02 | [−0.29, 0.25] | 346.79* |
| Gamma | 2 | 26 | 8 | −0.69 | [−3.00, 3.00] | 11.57 |
| Relative gamma | 3 | 24 | 7 | −1.09 | [−2.82, 0.63] | 7.16 |

*Note.* CI = confidence interval; *Q* = heterogeneity statistic; \* $p < .01$ .

### Subgroup Meta-analysis

#### Theta Electroencephalogram (Fig. 4A)

Theta band EEG data were subdivided based on EEG channels into frontal, parietal, and occipital channel subgroups for analyses. There were less than two studies regarding prefrontal, central, and temporal subgroups to conduct individual meta-analyses. We did not identify significant correlations with relaxation indices, and the omnibus test indicated no significant differences in the strength of correlations among channel subgroups (*QM*(2) = 0.13, *p* = .94). Although our study revealed significant heterogeneity for all channel subgroups (frontal: *Q* = 37.96, parietal: *Q* = 12.44, occipital: *Q* = 34.14), there were insufficient studies to conduct subgroup analysis.

**Figure 4.**
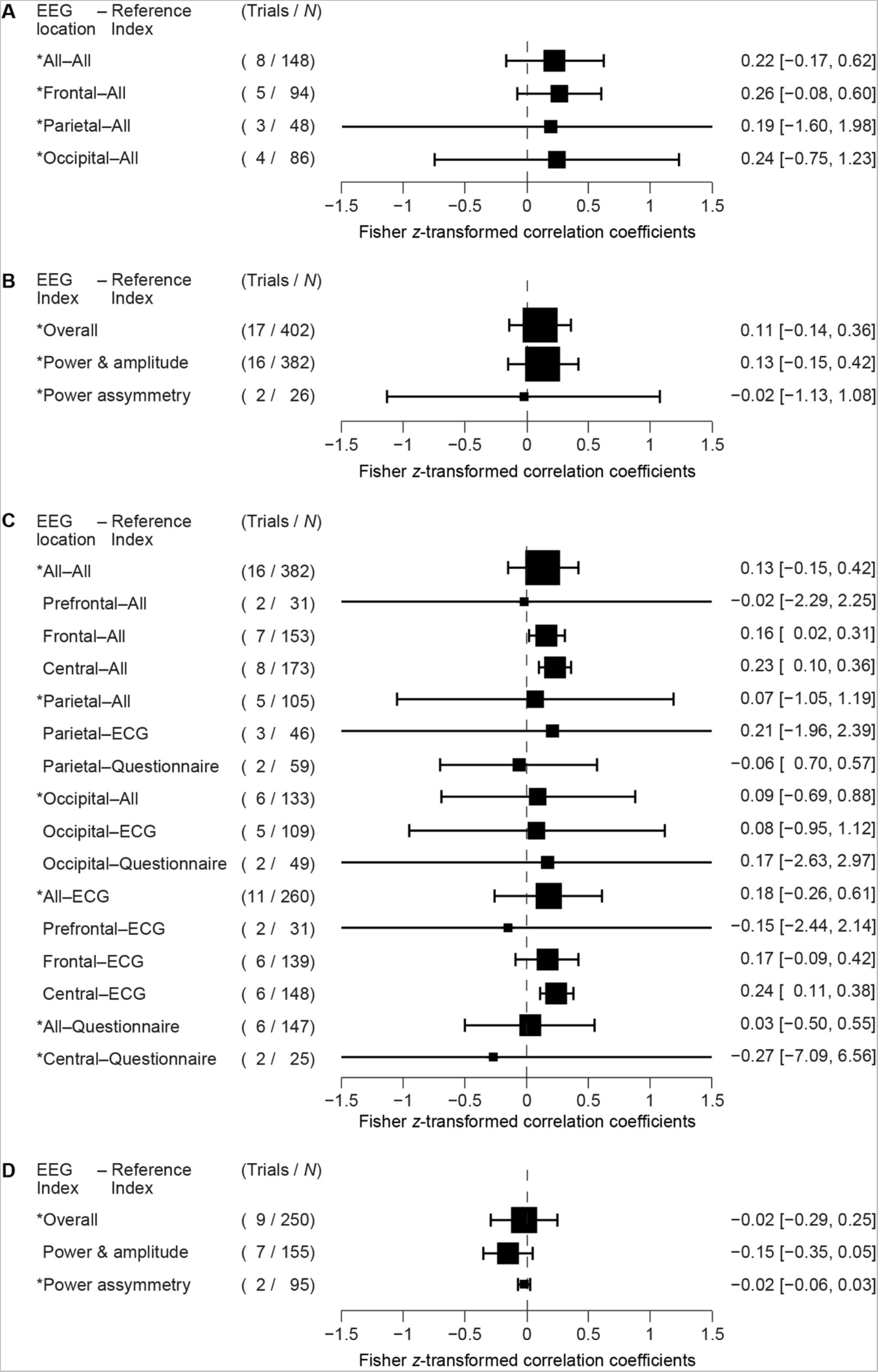
Forest plots for the correlation between electroencephalographic indices and reference indices for relaxation. (A) Correlation between theta power at different EEG sensor locations and reference indices for relaxation. (B) Correlation between different alpha indices and reference indices for relaxation. (C) Correlation between alpha power/amplitude at different EEG sensor locations and different reference indices for relaxation. (D) Correlation between different beta indices and reference indices for relaxation. Asterisks indicate that there was significant heterogeneity (*p* < .1 by Cochrane’s *Q* statistical test) among the trials. *N* = total number of subjects.

#### Alpha Electroencephalogram (Figs. 4B & 4C)

Alpha band data were subdivided to power/amplitude and power asymmetry, and both exhibited significant heterogeneity (power/amplitude: *Q* = 311.57, power asymmetry: *Q* = 51.35). Further alpha band power/amplitude were subdivided based on EEG channels into prefrontal, frontal, central, parietal, and occipital channel subgroups. Positive correlations were detected at frontal and central channels (frontal: *r* = 0.16, 95% CI [0.02, 0.31]; central: *r* = 0.23, 95% CI [0.10, 0.36]). Conversely, there were no correlations at prefrontal, parietal, and occipital channels. Consistent with these results, the omnibus test showed significant differences in correlation strength among channels (*QM*(4) = 9.99, *p* < .05), and post-hoc tests revealed that the correlation values were higher for the central subgroup than the other subgroups (compared with prefrontal and parietal: *p* < .01; compared to frontal and occipital: *p* < .05). No significant differences were found among the other channel subgroups. There was no heterogeneity in prefrontal, frontal, and central subgroups (prefrontal: *Q* = 9.07, frontal: *Q* = 30.30, central: *Q* = 46.00), but we observed heterogeneity in parietal and occipital subgroups (parietal: *Q* = 69.69, occipital: *Q* = 123.41). Further subgrouping within the parietal and occipital channels revealed that neither ECG indices nor questionnaire indices of relaxation were correlated with parietal or occipital alpha band power. Significant heterogeneity was not detected following this subgrouping, except for the occipital–ECG subgroup (parietal–ECG: *Q* = 35.22, parietal–questionnaire: *Q* = 13.43, occipital–ECG: *Q* = 120.04, occipital– questionnaire: *Q* = 3.36). We excluded the temporal channel subgroup from this analysis due to the limited number of outcome measures.

In addition to subgrouping based on EEG channels, alpha band data were subdivided based on reference indices into ECG-related and questionnaire subgroups, but no correlations were detected. Omnibus test showed significant differences in correlation strength between reference index types (*QM*(1) = 7.97, *p* < .01), and post-hoc tests revealed a significant difference between reference ECG-related indices or questionnaire subgroups (*p* < .001). Alpha–ECG-related indices and alpha–questionnaire subgroups exhibited significant heterogeneity (ECG: *Q* = 220.30, questionnaire: *Q* = 81.16). Further subgrouping was conducted for these subgroups, revealing significant correlation in the ECG–central subgroup (*r* = 0.24, 95% CI [0.11, 0.38]) but not in other combinations. The ECG–occipital and questionnaire–central subgroups exhibited significant heterogeneity (ECG–occipital: *Q* = 120.04, questionnaire–central: *Q* = 23.21), and other combinations exhibited no heterogeneity (ECG–prefrontal: *Q* = 3.76, ECG–frontal: *Q* = 26.22, ECG–central: *Q* = 22.36, ECG–parietal: *Q* = 35.22, questionnaire–parietal: *Q* = 13.43, questionnaire–occipital: *Q* = 3.36).

Subgroup analysis was not performed to investigate power asymmetry due to the limited number of outcome measures.

#### Beta Electroencephalogram (Fig. 4D)

Beta-band data were subdivided into power/amplitude and power asymmetry, but no correlations were detected. Power asymmetry subgroup exhibited significant heterogeneity (*Q* = 320.89), whereas power/amplitude subgroup exhibited no heterogeneity (*Q* = 18.57).

### Publication Bias

Distribution of correlation coefficients in the theta, alpha, and beta frequency categories were visualized using funnel plots (Fig. 5). Both tests suggested significant publication bias within the theta band (Egger’s test: *z* = −2.32, Begg’s test: τ = −0.57). Within the alpha band, significant publication bias was noted for Egger’s test (*z* = −1.73) but not for Begg’s test (τ = −0.26). In contrast, in the beta band, significant publication bias was observed for Begg’s test (τ = −0.67) but not for Egger’s test (*z* = −1.41). Due to the limited number of papers (≤3), we did not assess publication bias within the delta, gamma, and relative gamma frequency categories.

**Figure 5.**
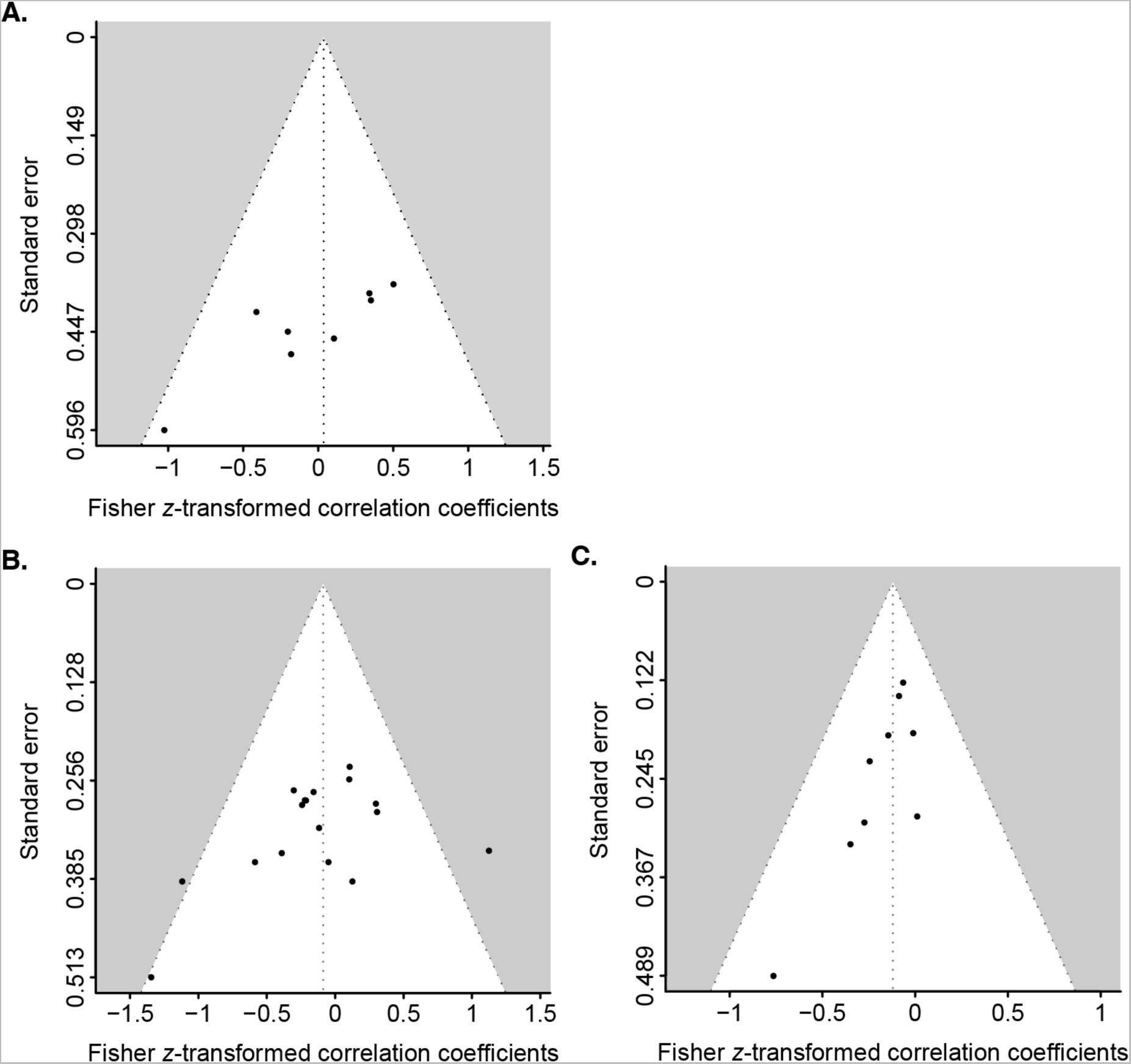
Funnel plots of the correlations between electroencephalographic indices and reference index of relaxation. (A) Correlation between theta indices and the reference index of relaxation. (B) Correlation between alpha indices and the reference index of relaxation. (C) Correlation between beta indices and the reference index of relaxation. Solid circles represent effect sizes.

## Discussion

The association between relaxation and alpha power is well-documented, but previous meta-analyses have focused primarily on a specific type of relaxation method (Bolouki, 2023; Han et al., 2022; Komini et al., 2023; Lomas et al., 2015; Wahbeh et al., 2018; Zaccaro et al., 2018) or a specific population (Jackson et al., 2015; Thabrew et al., 2018; Thibodeau et al., 2006). Further, most previous systematic reviews did not conduct meta-analyses. The current meta-analysis is distinguished by examination of the general association between EEG metrics and relaxation among healthy adults using data pooled from studies reporting many different indices and relaxation methods (Tables 1 and 2). From the 3,295 academic references, 38 studies were chosen for qualitative synthesis, and 23 of these were included in the meta-analysis. The results revealed a significant positive correlation between relaxation and the power/amplitude of alpha oscillations at central and frontal electrode channels. Furthermore, the correlation at central channel was significantly stronger than those detected at other channel locations. These results thus suggest that alpha power at central electrode may serve as a good indicator of relaxation in healthy adults.

This study confirms the importance of electrode location when examining the link between alpha oscillations and relaxation. Specifically, our findings suggest that central and frontal alpha oscillations and their underlying neural processes are associated with relaxation. Although central and frontal electrode measures were significantly correlated with relaxation, other electrodes yielded no significant correlations. Occipital alpha oscillations are more pronounced when individuals are at rest with their eyes closed. These occipital oscillations have been suggested to originate from interactions between the thalamus and occipital cortices (Minami et al., 2020) and are closely associated with perception and cognition (Amano et al., 2008; Busch et al., 2009; Kawashima et al., 2023; Mathewson et al., 2009; Minami & Amano, 2017; Samaha & Postle, 2015; van Dijk et al., 2008). Conversely, alpha oscillations in the central channel—also known as Rolandic rhythm (Gastaut et al., 1954)—are known to decrease during motor planning, reflecting cognitive and psychomotor load (Bazanova & Vernon, 2014; Erbil & Ungan, 2007). As relaxation involves intentional inhibition of motor planning, the cortical motor system may subserve the relationship between central alpha oscillations and relaxation.

Conversely, alpha oscillations in the frontal cortex have been suggested to reflect the control of attention (Aftanas & Golocheikine, 2001; Magosso et al., 2021; Misselhorn et al., 2019) and contribute to more efficient neurocognitive processing (Berger & Davelaar, 2018). Therefore, the relationship between relaxation and alpha oscillations in frontal channels may be mediated by attention. Alternatively, considering the involvement of the thalamus—which also interacts with alpha oscillations—in ANS regulation (Cechetto, 2014; Kuriyama et al., 2010; Morrison, 2001; Rachão et al., 2021; Halgren et al., 2019; Hughes & Crunelli, 2005), thalamocortical interactions may play a role in the relationship between alpha oscillations and relaxation. Moreover, these relationships may be even stronger if the alpha oscillation band is further divided into narrower frequency sub-bands, as seven out of the 38 studies in the qualitative summary subdivided alpha oscillations into low- and high-frequency ranges. This finer analysis is also important as different sub-bands may reflect distinct functional aspects of relaxation (Bazanova & Vernon, 2014; Klimesch et al., 1998; Shibusawa et al., 2023).

It should be noted that the present meta-analysis assessed only the relationship between power/amplitude and relaxation state, due to the limited number of studies utilizing other EEG indices. For example, while one study reported a negative correlation between alpha-peak frequency at left occipital channels and calm component score (Kabuto et al., 1993), this data could not be included in the current meta-analysis as there were no other similar studies. Furthermore, we could not investigate the association between relaxation and frontal alpha asymmetry, known to be associated with depression and negative mood (Davidson, 1993; Yadollahpour et al., 2019), due to an insufficient number of studies. To draw more solid conclusions about the association between EEG oscillations and relaxation, further research is needed using a variety of EEG indices.

The causal relationship between alpha oscillations and relaxation remains unclear, although a few studies have explored the potential for alpha oscillation neurofeedback to induce relaxation (Biswas & Ray, 2019; Grosselin et al., 2021). However, it is also possible that the change in alpha power is induced by relaxation. To examine the causal association between alpha oscillations and relaxation, it is necessary to investigate whether relaxation states can be induced by entraining alpha oscillations, such as by using brain stimulation methods like transcranial magnetic stimulation or transcranial alternating current stimulation.

An important limitation of this meta-analysis is that there is yet no established standard index for relaxation state. In this meta-analysis, reference indices included both introspective and physiological measures linked to the ANS. However, relaxation is inherently complex, and psychological differences among individuals may influence subjective ratings of relaxation. More critically, introspective assessments of relaxation may actually influence neural activity (Block, 2019; Tsuchiya et al., 2015). It is also unclear whether ANS-related indices accurately predict introspective relaxation (LeDoux & Brown, 2017; LeDoux & Pine, 2016).

No relationship with relaxation was found in EEG frequency bands other than alpha. Thus, the current meta-analysis does not support the relationships between theta oscillations and relaxation reported in previous studies (Chan et al., 2008; Lomas et al., 2015; Sinha et al., 2020). In addition, due to a limited number of studies, associations of relaxation with gamma oscillations, delta oscillations, and power ratios between frequencies remain uncertain. Finally, risk of bias assessment revealed a noticeable absence of studies with pre-registration and blinding of outcome assessors, underscoring the need for more rigorous and objective research practices for examining these associations.

In summary, the current systematic review and meta-analysis revealed specific positive correlations between relaxation and the power of alpha oscillations at central and frontal EEG channels. Furthermore, the correlation coefficient for the central channel was stronger than that for other channels. However, due to limited investigation beyond alpha power/amplitude, much further research is needed to comprehensively describe the EEG correlates of relaxation.

## Acknowledgments

We thank the members of the Neurotech Evidence Evaluation Committee, a specially organized group for the JST Moonshot R&D project, for their comments on the manuscript. A special thanks also go to Dr. Ryoji Onagawa and Prof. Katsumi Watanabe for their support.

References marked with an asterisk indicate studies included in the meta-analysis.

